# The making of calibration sausage exemplified by recalibrating the transcriptomic timetree of jawed vertebrates

**DOI:** 10.1101/2019.12.19.882829

**Authors:** David Marjanović

## Abstract

Molecular divergence dating has the potential to overcome the incompleteness of the fossil record in inferring when cladogenetic events (splits, divergences) happened, but needs to be calibrated by the fossil record. Ideally but unrealistically, this would require practitioners to be specialists in molecular evolution, in the phylogeny and the fossil record of all sampled taxa, and in the chronostratigraphy of the sites the fossils were found in. Paleontologists have therefore tried to help by publishing compendia of recommended calibrations, and molecular biologists unfamiliar with the fossil record have made heavy use of such works (in addition to using scattered primary sources and copying from each other). Using a recent example of a large node-dated timetree inferred from molecular data, I reevaluate all thirty calibrations in detail, present the current state of knowledge on them with its various uncertainties, rerun the dating analysis, and conclude that calibration dates cannot be taken from published compendia or other secondary or tertiary sources without risking strong distortions to the results, because all such sources become outdated faster than they are published: 50 of the sources I cite to constrain calibrations were published in 2019, half of the total of 276 after mid-2016, and 90% after mid-2005. It follows that the present work cannot serve as such a compendium either; in the slightly longer term, it can only highlight known and overlooked problems. Future authors will need to solve each of these problems anew through a thorough search of the primary paleobiological and chronostratigraphic literature on each calibration date every time they infer a new timetree; and that literature is not optimized for that task, but largely has other objectives.

## 1 Introduction

This work is not intended as a review of the theory or practice of node (or tip) dating with calibration dates (or tip dates) inferred from the fossil record; as the most recent reviews of methods and sources of error I recommend those by Barido-Sottani et al. (2019, 2020), Matschiner (2019), Marshall (2019), Guindon (2020), Powell et al. (2020), Pardo et al. (2020), and, with caveats of which I will address two (Materials and methods: Calibrations: Node 152 – Placentalia), Springer et al. (2019). Neither is it intended as a review of the history of the dates assigned to certain calibrations; as an example of a recent detailed review of three commonly used calibrations, I recommend Pardo et al. (2020). Although I discuss wider implications, the scope of this work is narrow: to evaluate each of the 30 calibrations used in the largest vertebrate timetree yet published, that by Irisarri et al. (2017), and the total impact of the errors therein on the results (using the same node-dating method they used, which I do not evaluate beyond mentioning potential general points of criticism).

Irisarri et al. (2017) inferred a set of timetrees from the transcriptomes of 100 species of gnathostomes (jawed vertebrates) and combinations of up to 30 calibrations from the fossil record. On the unnumbered ninth page of their supplementary information, they described their calibration dates as “five well-accepted fossil calibrations plus a prior on the root” and “24 additional well-established calibration points with solid paleontological evidence”. For many of the calibrations, these optimistic assessments are not tenable. I have tried to present, and use, the current state of knowledge on each of these calibrations.

In doing so, the present work naturally resembles the compendia of suggested calibrations that paleontologists have occasionally compiled with the intent to provide a handy reference for molecular biologists who wish to date divergences (e.g. Müller and Reisz, 2007; Benton et al., 2015, and six other articles in *Palaeontologia Electronica* 18(1); Wolfe et al., 2016; Morris et al., 2018); Irisarri et al. (2017) took seven of their 30 calibrations from the compendium in Benton and Donoghue (2007: table 1) alone – without citing the enlarged update by Benton et al. (2015) –, compared to six taken from the primary literature. However, I will show that all such compendia are doomed to be (partially) outdated almost as fast as they are published in the best case, and faster than they are published in the average case. Soon, therefore, the present work will no longer be reliable as such a compendium either; rather, it is intended to show readers where the known uncertainties and disagreements lie, and thus what anybody who wants to use a particular calibration should probably search the most recent literature for. This is why I do not generally begin my discussion of a calibration by presenting my conclusions on what the best, or least bad, minimum and maximum ages of the calibration may be. (They are, however, presented without further ornament in Table 1.) Instead, I walk the reader through a sometimes meandering discovery process, demonstrating how this knowledge was arrived at and how it may soon change – how the sausage was made and how it may spoil.

**Table 1:**
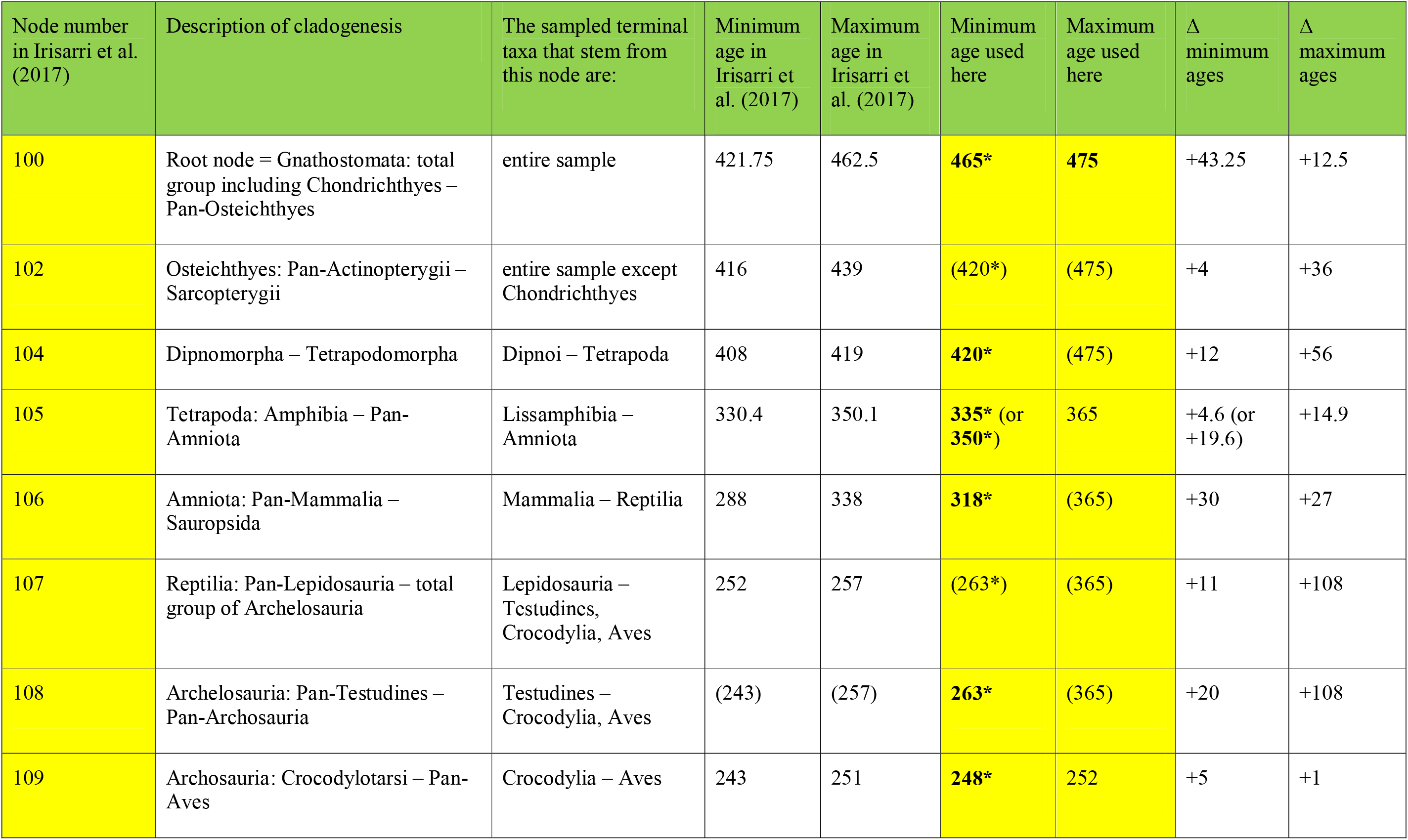
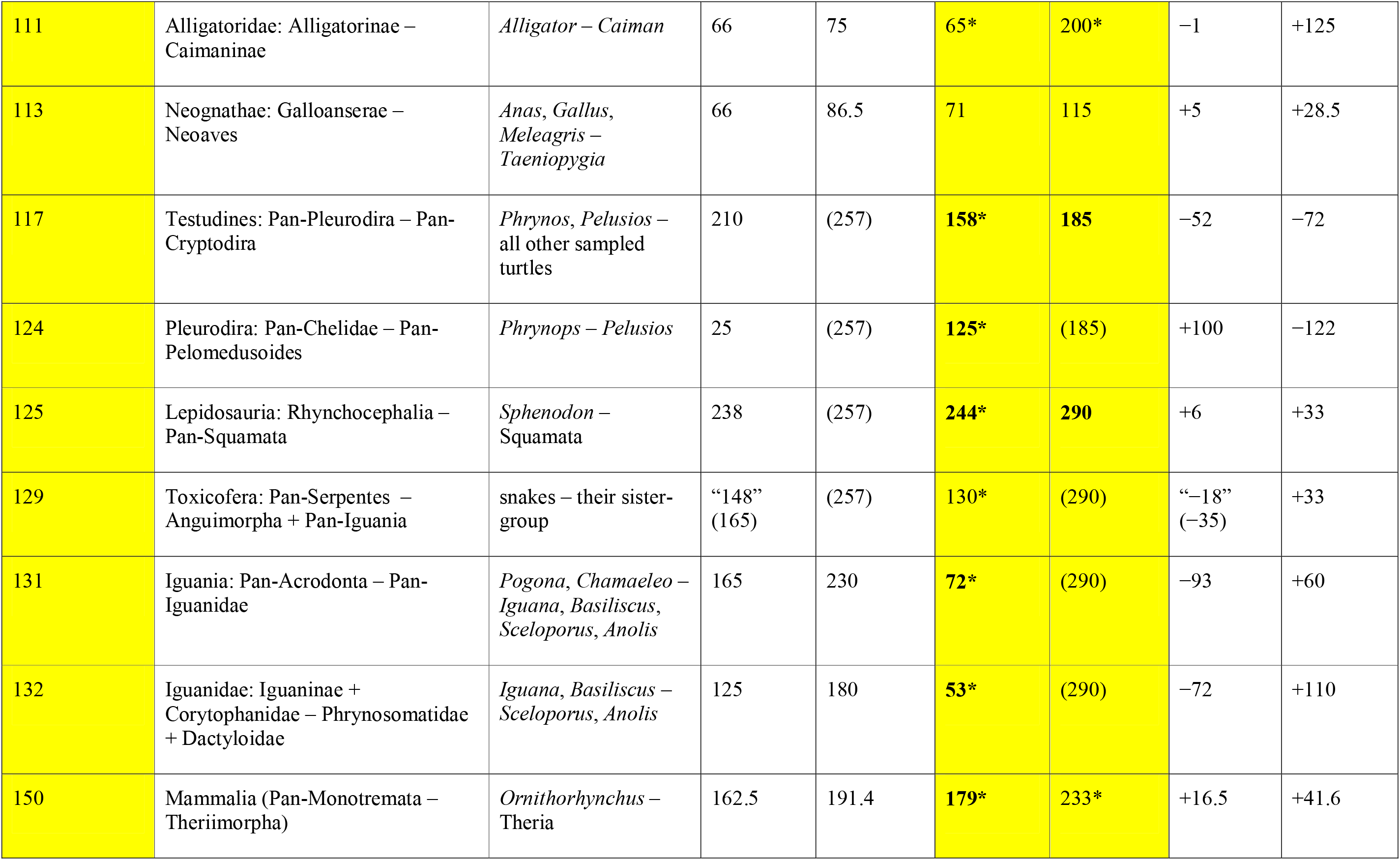
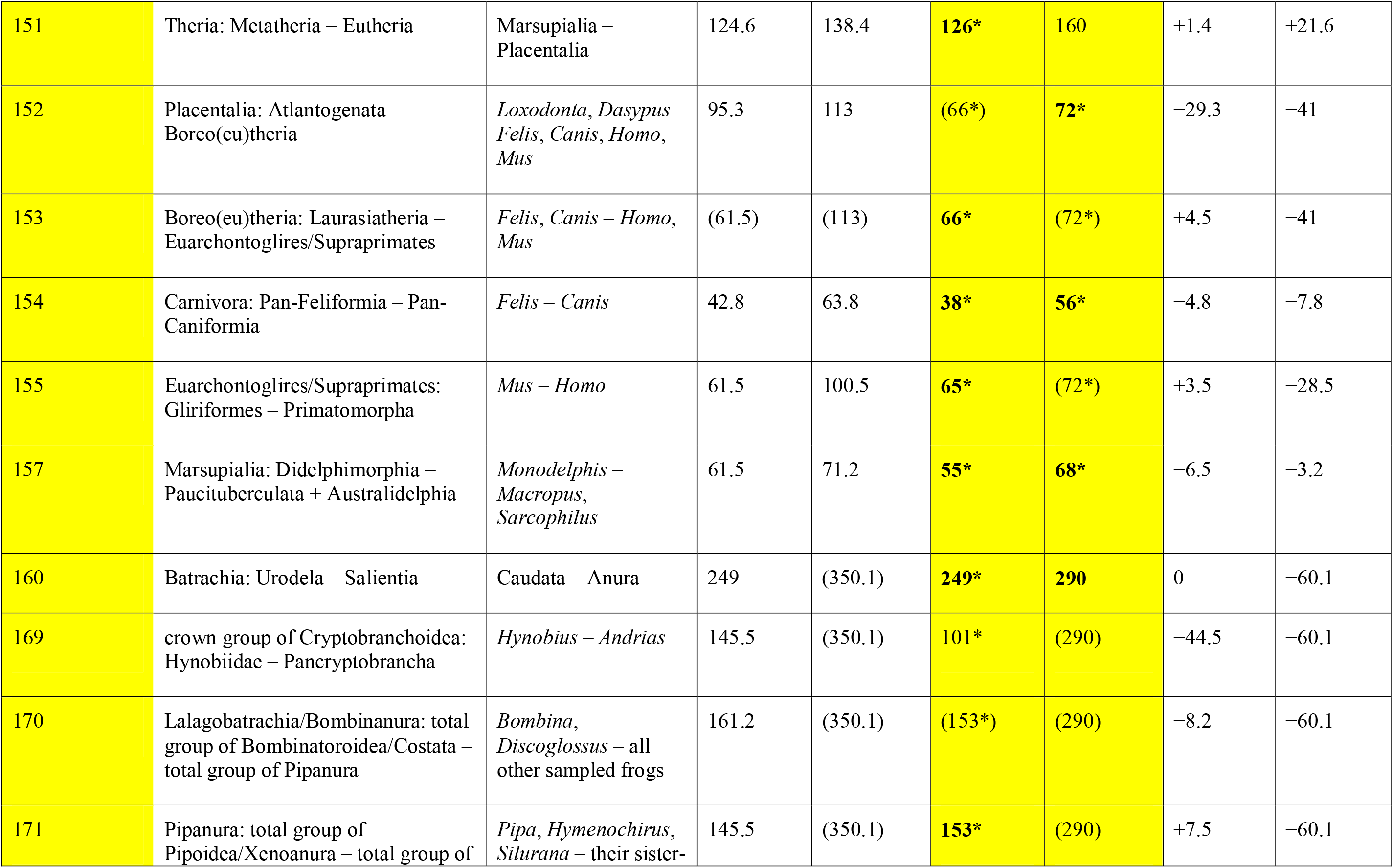
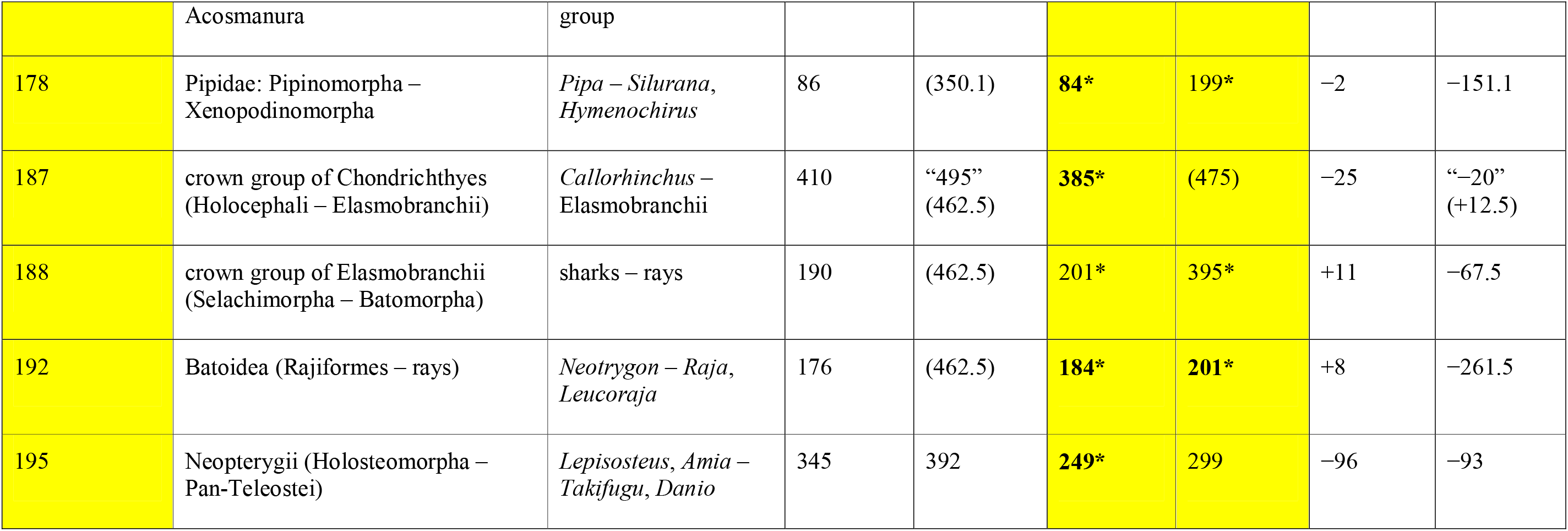
The first four columns of Irisarri et al. (2017: supplementary table 8 and supplementary figure 19), here expanded to five, followed by the ages used here for the same calibrations and the differences (Δ). Boldface is a rough indicator of my confidence. Hard bounds are marked with an asterisk. Dates in parentheses were not specified in the analysis; the node was constrained in practice by the given constraint on a preceding (for maximum ages) or following node (for minimum ages) elsewhere in this table – see Fig. 1 for which nodes precede each other. The two dates in quotation marks were specified by Irisarri et al. (2017), but had no effect because they were in practice constrained by the dates specified for other nodes. Dashes in the second and third column separate the two branches stemming from the node in question. Depending on the node, see the text or the Supplementary Material for discussion and references.

Some works used as compendia in this sense are not even compiled by paleontologists: molecular biologists often copy from each other. Irisarri et al. (2017) took four of their calibrations from table 1 of Noonan and Chippindale (2006), a work that contains a phylogenetic and divergence-date analysis of molecular data and cites severely outdated paleontological primary and secondary literature (from 1981 to 2003) as its sources.

A continually updated online compendium could largely avoid the problem that knowledge has a half-life. There has been one attempt to create one, the Fossil Calibration Database (Ksepka et al., 2015 – https://fossilcalibrations.org; not counting separately its predecessor, called Date a Clade, which is no longer online and apparently merely presented table 1 of Benton and Donoghue, 2007). It appears to have run out of funding long ago and has not been updated since 2 February 2018, the day on which three of the numerous calibrations proposed in Wolfe et al. (2016) were added; other calibrations from the same source were added on 30 and 31 January 2018 (one each) and 22 December 2017 (three), and no other updates were made on those days. I cannot resist pointing out that this is one of many cases where funding menial labor in the sciences – reading and interpreting papers, evaluating the contradictions between them, and entering the interpretations in a database, a task that cannot be automated – would go a long way toward improving the quality of a large number of publications, but is unlikely to be granted because it is not likely to result in a single flashy publication or in an immediately marketable application directly, even though precise and accurate timetrees are an essential component of our understanding of the model organisms used in biomedical research.

A continually updated online database aiming to represent the entire fossil record exists, and is currently being funded: the Paleobiology Database, accessible through two different interfaces at http://www.pbdb.org and https://paleobiodb.org. Among many other things, it aims to contain the oldest currently known record of every taxon and would thus be useful as a source for calibrations. However, the warnings by Parham et al. (2011) still apply: the quality of the Paleobiology Database is quite heterogeneous. While some entries are written by the current top experts in the respective fields, others copy decades-old primary descriptive literature uncritically, often leading to severely outdated taxonomic, let alone phylogenetic placements (in all but the most recent literature that is not the same), not to mention misunderstandings based on the convoluted history of taxonomic nomenclature. It is not uncommon for two entries to contradict each other. Finally, despite the hundreds of contributors, our current knowledge of the fossil record is so vast that the database remains incomplete (again, of course, differently so for different taxa). Like Irisarri et al. (2017), I have not used the Paleobiology Database or the Fossil Calibration Database; I have relied on the primary literature.

### 1.1 Nomenclature

After the publication of the *International Code of Phylogenetic Nomenclature (PhyloCode)* (Cantino and de Queiroz, 2020) and its companion volume *Phylonyms* (de Queiroz et al., 2020), the registration database for phylogenetic nomenclature – *RegNum* (Cellinese and Dell, 2020) – went online on 8 June 2020; regulated phylogenetic nomenclature is therefore operational. In an effort to promote uniformity and stability in nomenclature, I have used the names and definitions from *Phylonyms* and Ezcurra et al. (2020: online methods) here; wherever applicable, all of them are followed by “[PN]” at least at the first mention (this includes vernacularized forms like “gnathostome”) to avoid confusion with earlier uses of the same names for different clades. I have not, however, followed the *ICPN*’s Recommendation 6.1A to set all taxonomic names in italics.

The definitions of these names, their registration numbers (which establish priority among the combinations of name and definition) and the exact chapter citations can be found in *RegNum*, which is freely accessible (https://www.phyloregnum.org/).

*ICPN*-regulated names have not been created or converted according to a single overarching scheme. As a result, for example, the name Osteichthyes has been defined as applying to a crown group, and the corresponding total group has been named Pan-Osteichthyes; but the name Chondrichthyes has not been defined and could end up as the name for a crown group, a total group, or neither (indeed, current common usage by paleontologists is neither). This has required some awkward circumlocutions. Following Recommendation 9B of the *ICPN*, I have not coined any new names or definitions in the present work.

The shapes and definitions of most other taxonomic names used here do not currently compete for homonymy or synonymy under any code of nomenclature. (The *ICPN* is not retroactive, and the rank-based *International Code of Zoological Nomenclature* [ICZN, 1999] does not regulate the priority of names at ranks above the family group.) In such cases, I have followed current usage where that is trivial; I occasionally mention synonyms where that seems necessary.

The usage of “stem” and “crown” requires a comment. The crown group of a clade consists of the last common ancestor of all extant members of that clade, plus all its descendants. The rest of the clade in question is its stem group. For example, *Gallus* is a crown-group dinosaur, and *Triceratops* is a stem-group dinosaur. In a development that seems not to have been foreseen by the first two or so generations of phylogeneticists that established the terminology – for example, the zoology textbook by Ax (1987) exclusively named total groups, i.e. halves of crown groups! –, many clades with defined names are now identical to their crown groups (in other words, they are crown clades); they do not contain any part of their stem. Aves [PN] is an example; although *Triceratops* is a stem-dinosaur [PN], a stem-dinosauromorph [PN] and a stem-ornithodiran [PN] among other things, it is not a stem-bird or stem-avian because by definition there is no such thing. It is instead a stem-pan-avian [PN], i.e. a stem-group member of Pan-Aves [PN] (Ezcurra et al., 2020: online methods). If no name is available for a suitable larger group, I have resorted to the circumlocution that *Triceratops*, for instance, is “on the bird stem” or “in the avian total group” (expressing that it is closer to Aves than to any mutually exclusive crown group).

## 2 Materials and methods

Although I have followed the spirit of the guidelines developed by Parham et al. (2011) for how best to justify or evaluate a proposed calibration, I have not consistently followed their letter. Most notably, the specimen numbers of the fossils that I largely refer to by genus names can all be found in the directly cited primary literature, so they are not repeated here.

### 2.1 Hard and soft minima and maxima

Without discussing the matter, Irisarri et al. (2017) stated that they had treated all calibration ages as soft bounds, which, in the software they used, means that “a proportion of 0.05 of the total probability mass is allocated outside the specified bound(s) (which means, 5% on one side, in the cases of the pure lower and pure upper bounds, and 2.5% on each side in the case of a combination of lower and upper bound)” (Lartillot, 2015: manual). This is particularly odd for minimum ages; after all, the probability that a clade is younger than its oldest fossil is not 5% or 2.5%, it is 0%. A few other works have used soft minima as an attempt to account for phylogenetic or chronostratigraphic uncertainty of the specimens chosen as calibrations. I have not used the former approach here (despite two clumsy attempts in the first preprint of this paper – Marjanović pointed out as incoherent by a reviewer): in the cases of phylogenetic uncertainty discussed below, different fossils that could calibrate the age of a cladogenetic event are commonly tens of millions of years apart, a situation that cannot be smoothed over by using the oldest one as a soft minimum. Soft minima that can be justified by uncertainty over the exact age of a calibrating fossil are very rare nowadays (as already pointed out by Parham et al., 2011); within the scope of this paper there is only one such case, the minimum age of Neognathae (node 113), which is determined by a specimen that is roughly 70 ± 1 Ma old according to a fairly long chain of inference. I have treated all other minima as hard, and I have not spelled this out below.

As recommended by Parham et al. (2011), minimum ages have generally been chosen in the literature as the youngest possible age of the calibrating specimen(s). This is practically guaranteed to result in ages that are too young for various reasons (Marshall, 2019). To account, if crudely, for non-zero branch lengths and especially for the nested phylogenetic positions of some calibrating specimens, and to counteract “the illusion of precision” (Graur and Martin, 2004: title) spread by calibration ages with five significant digits like 421.75 Ma (the minimum age chosen by Irisarri et al. [2017] for the root node, see below), I have rounded up (stratigraphically down) to the nearest million years, with a few exceptions suggested by mass extinction events.

Maximum ages are by default much more difficult to assign than minimum ages. Absence of proof is not proof of absence; absence of evidence is evidence of absence, but in most cases it is quite weak evidence. Yet, omitting maximum ages altogether and assigning only minimum ages to all calibrations automatically results in much too old divergence dates as nothing stops the 99.9% or 99.99% confidence or credibility intervals for all node ages from avoiding all overlap with the calibrated minimum ages. I have therefore followed Irisarri et al. (2017) and their sources in assigning as many maximum ages as I dare. For this purpose I have basically followed the recommendations of Parham et al. (2011) and Pardo et al. (2020: 11), which amount to assigning a maximum age whenever we can reasonably expect (after preservation biases, collection biases, collection intensity, paleobiogeography etc.) to have found evidence of the clade in question if it had been present at the time in question, but have not found any. This has widely been followed in the literature, but various compendia like Benton et al. (2015) have gone beyond this in many cases: in short, the oldest certain fossil provides the minimum age under that approach, while the oldest uncertain fossil of the same clade provides the maximum age. This practice is not defensible; therefore I assign, in the aggregate, fewer and more distant maximum ages than Irisarri et al. (2017).

Given the limits of our current knowledge of the fossil record, all maximum ages might be expected to be soft bounds. In a few cases discussed below, however, I find that the absence of evidence is so hard to explain away that a hard maximum is justified. This generally concerns unrealistically old maxima that I have chosen because no younger maximum suggests itself. Ultimately, of course, this is subjective.

The choices of hard vs. soft bounds do not seem to make a great difference to the big picture. Due to practical constraints, a set of calibration ages mostly identical to the present ones was analyzed twice, with all bounds treated as soft or as hard, in the first preprint of this work (Marjanović, 2019); the results were quite similar to each other (Marjanović, 2019: fig. 1, table 2). Even so, however, in the run where all bounds were soft, most divergence dates were younger than in the run where all bounds were hard (usually negligibly so, but by 20 Ma in the extreme cases); the mean ages of some calibrated nodes even ended up younger than their minimum ages.

**Figure 1:**
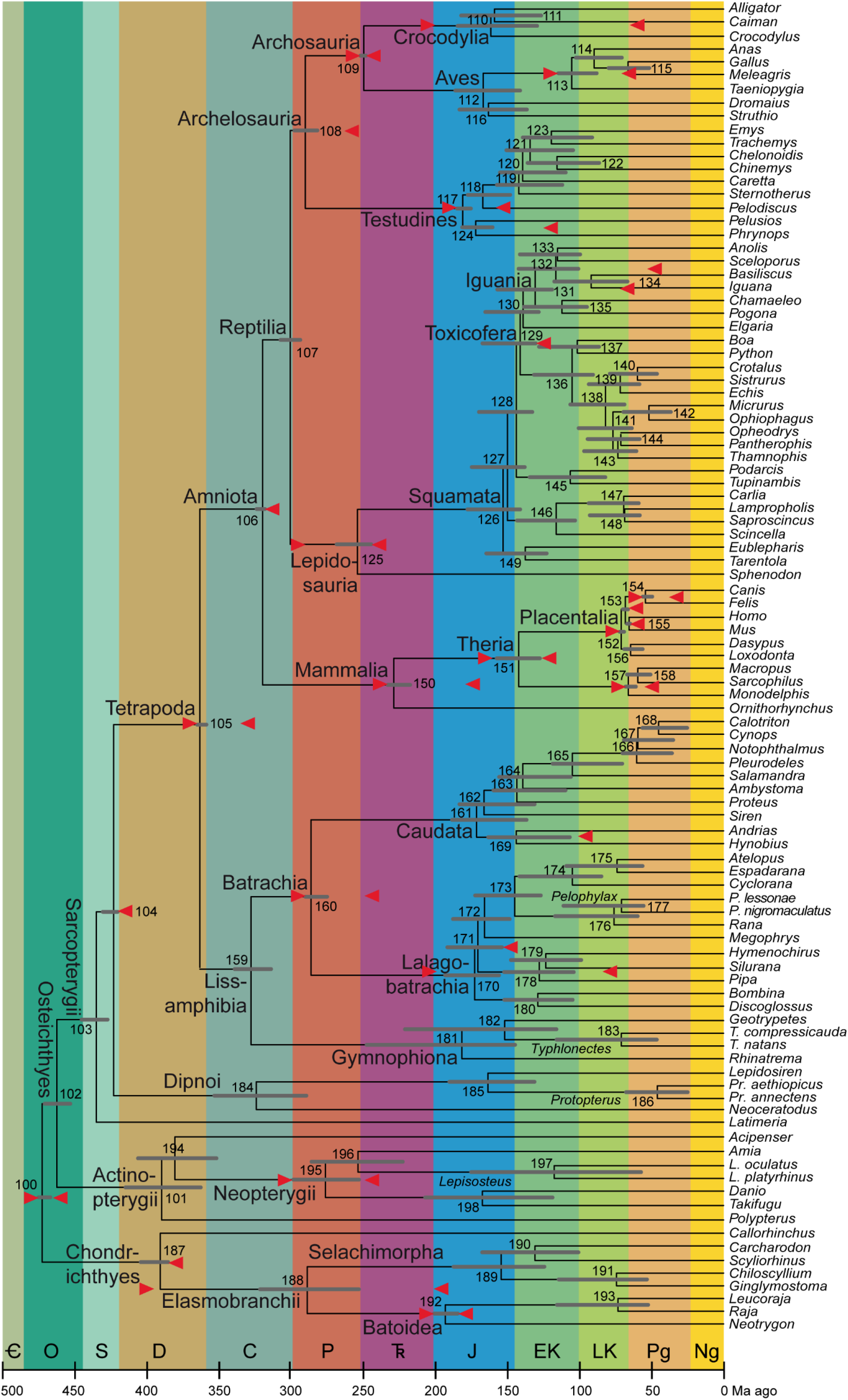
Average timetree resulting from application of the calibrations described here. As in Table 2 and in Irisarri et al. (2017: fig. 3), the bars on the nodes are the superimposed 95% credibility intervals from the two runs in PhyloBayes. The calibrations are shown as red arrows horizontally in line with the nodes they apply to; note that the arrow that is almost aligned with the branch of Lalagobatrachia and the one that is almost aligned with the terminal branch for *Silurana* are the maximum and minimum ages of node 178 (Pipidae), the one on *Iguana* to node 131 (Iguania), and the one on *Pelodiscus* to node 117 (Testudines). The abbreviated genus names are spelled out as clade names on their common branches; where only one species per genus is sampled, see Irisarri et al. (2017) for full species names. To the extent possible, clade names with minimum-clade (node-based) definitions are placed close to those nodes, while names with maximum-clade (branch-based) definitions are shown close to the origin of that branch (i.e. the preceding node if sampled) and undefined names stay in the middle. Period/epoch symbols from oldest to youngest: Cambrian (cut off at 500 Ma), Ordovician, Silurian, Devonian, Carboniferous, Permian, Triassic, Jurassic, Early Cretaceous, Late Cretaceous, Paleogene, Neogene including Quaternary (which comprises the last 2.58 Ma and is not shown separately). Timescale (including colors) from the International Chronostratigraphic Chart, version 2020/03 (Cohen et al., 2020). Node numbers, also used in the text and the Tables, from Irisarri et al. (2017).

**Table 2:**
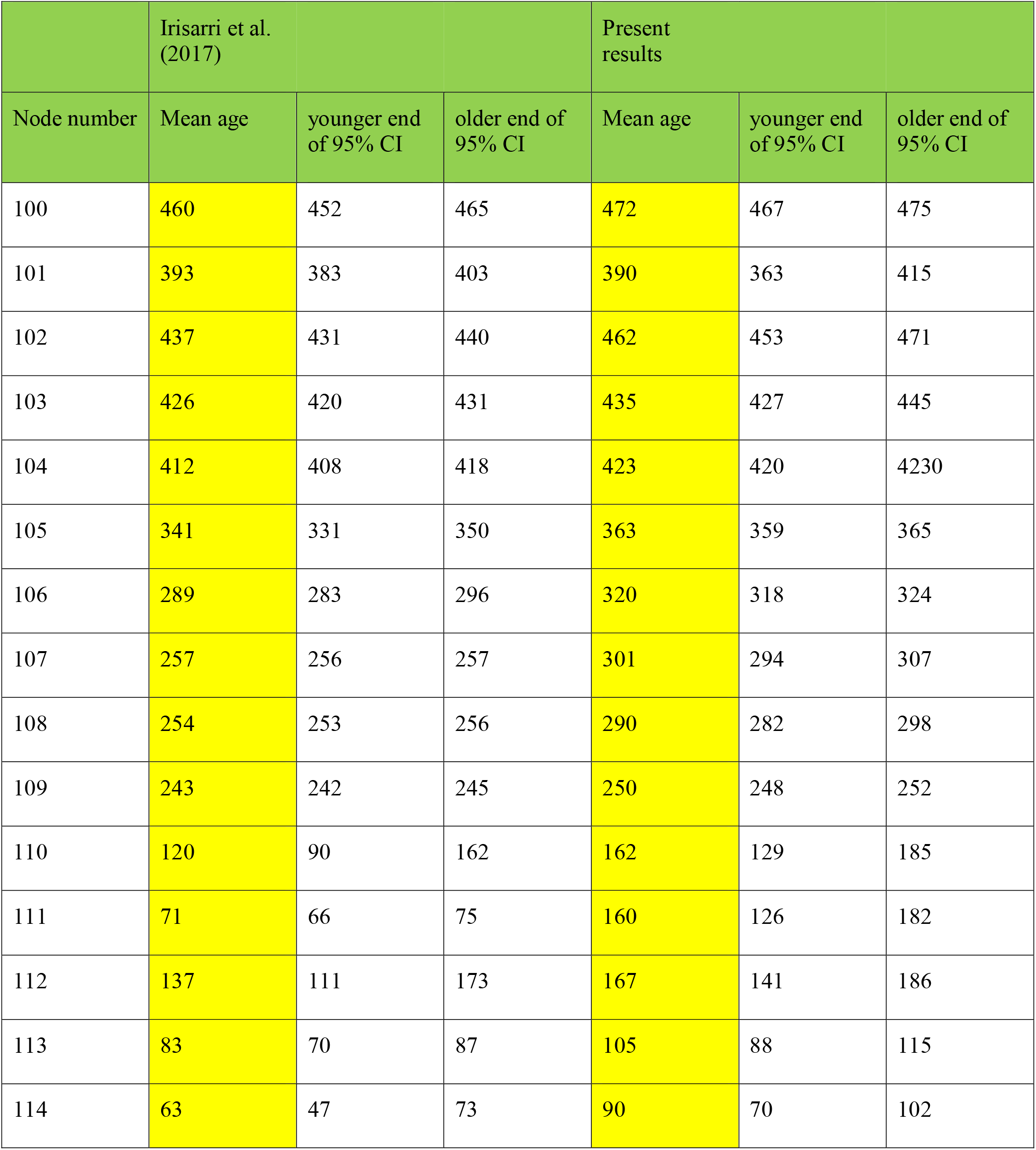
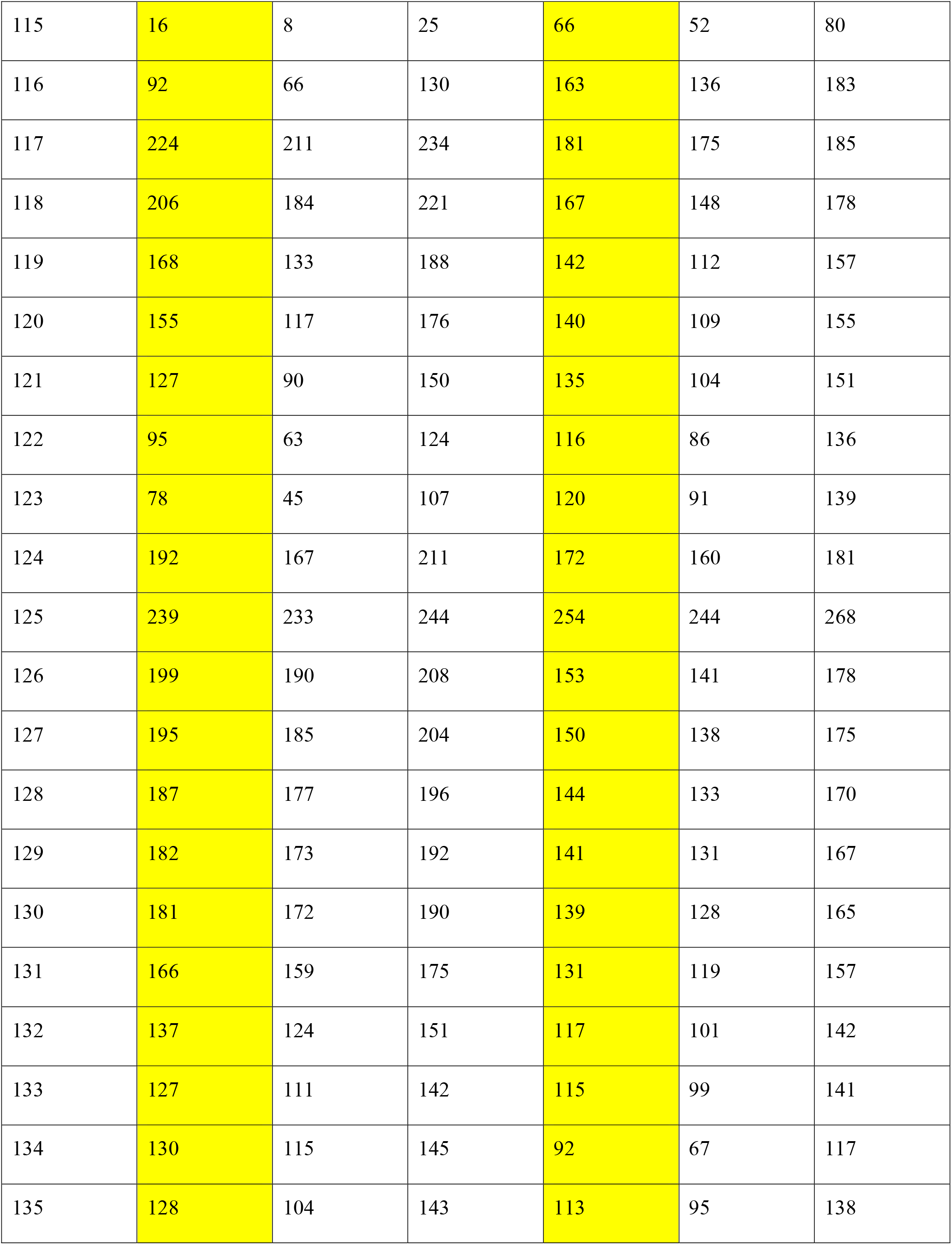
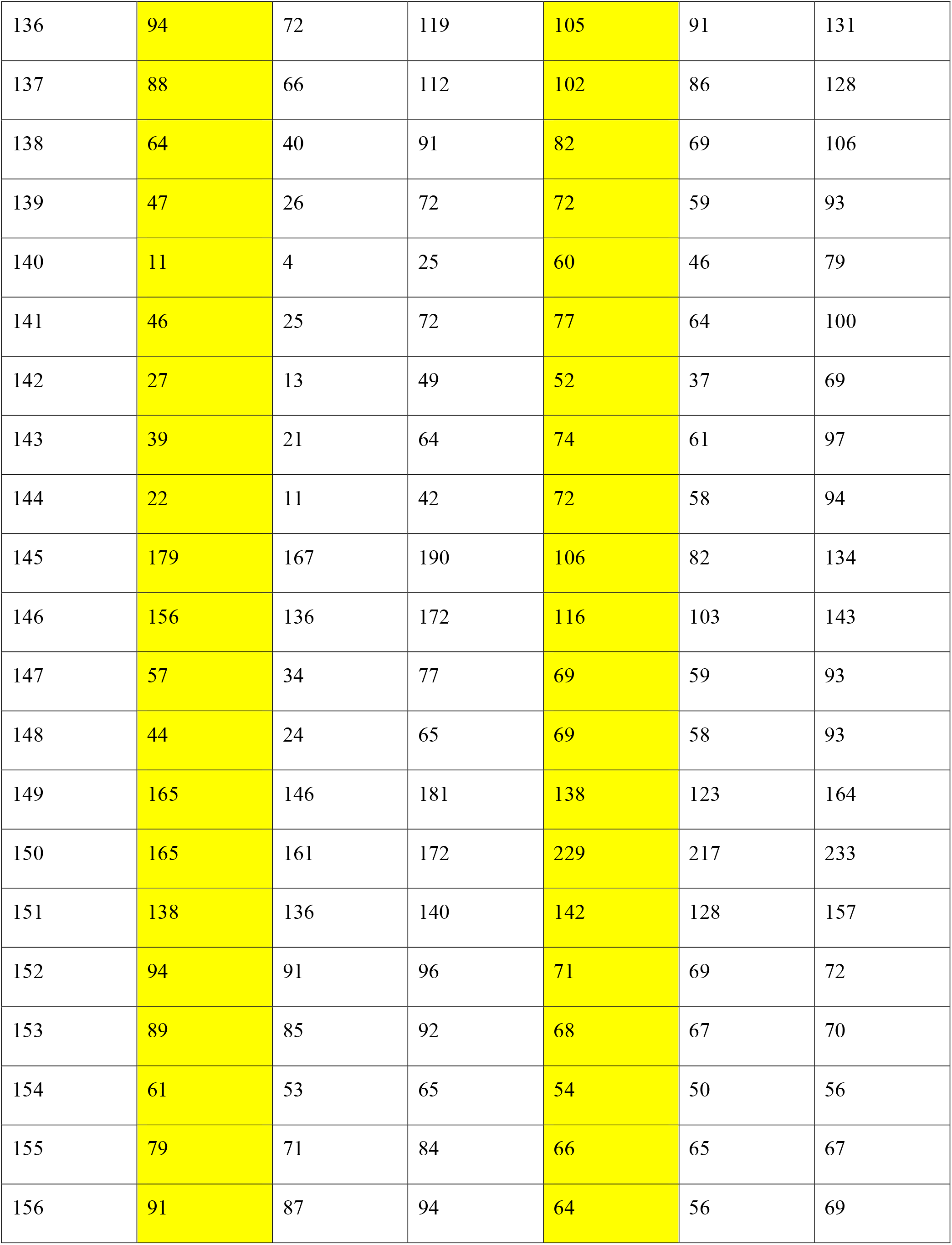
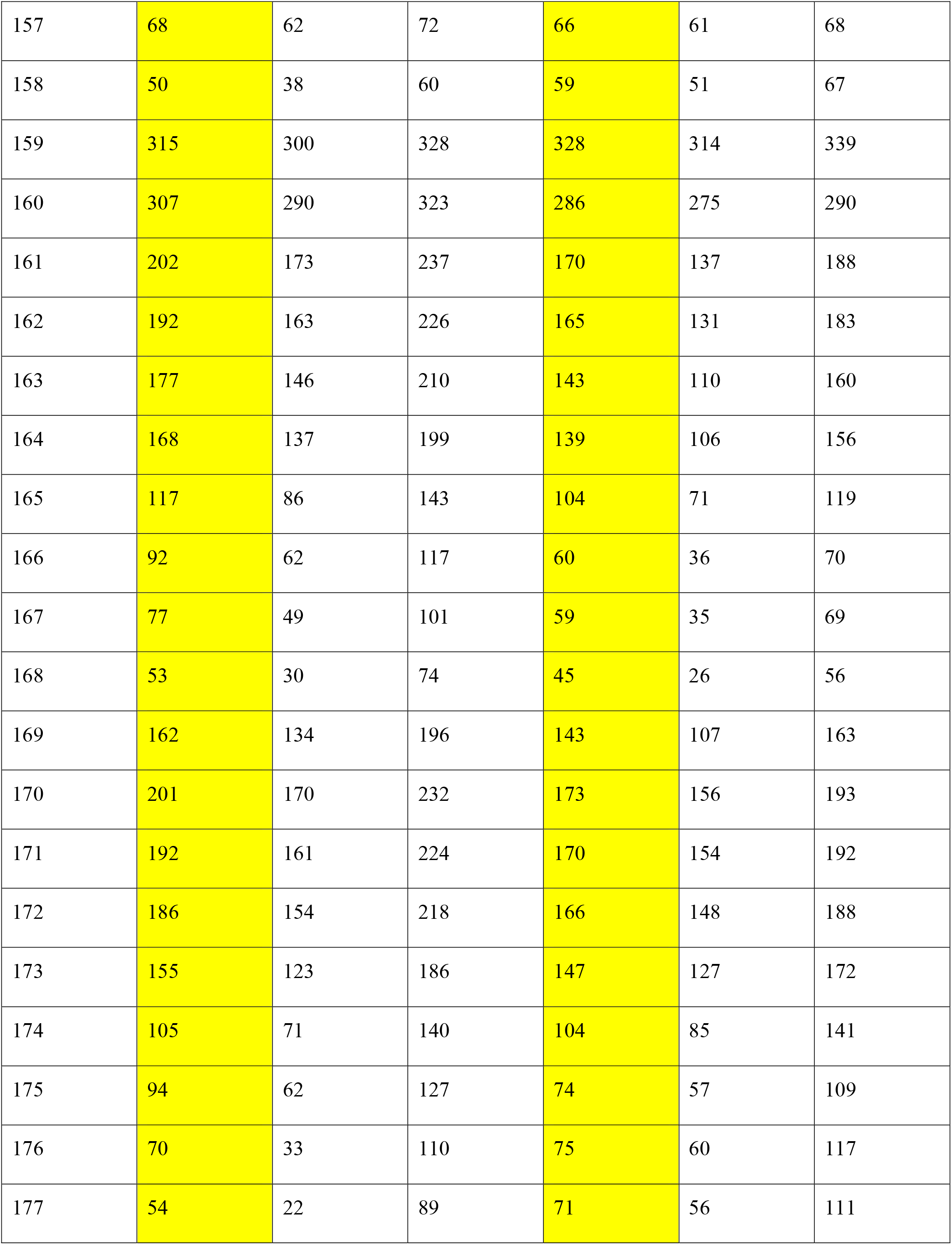
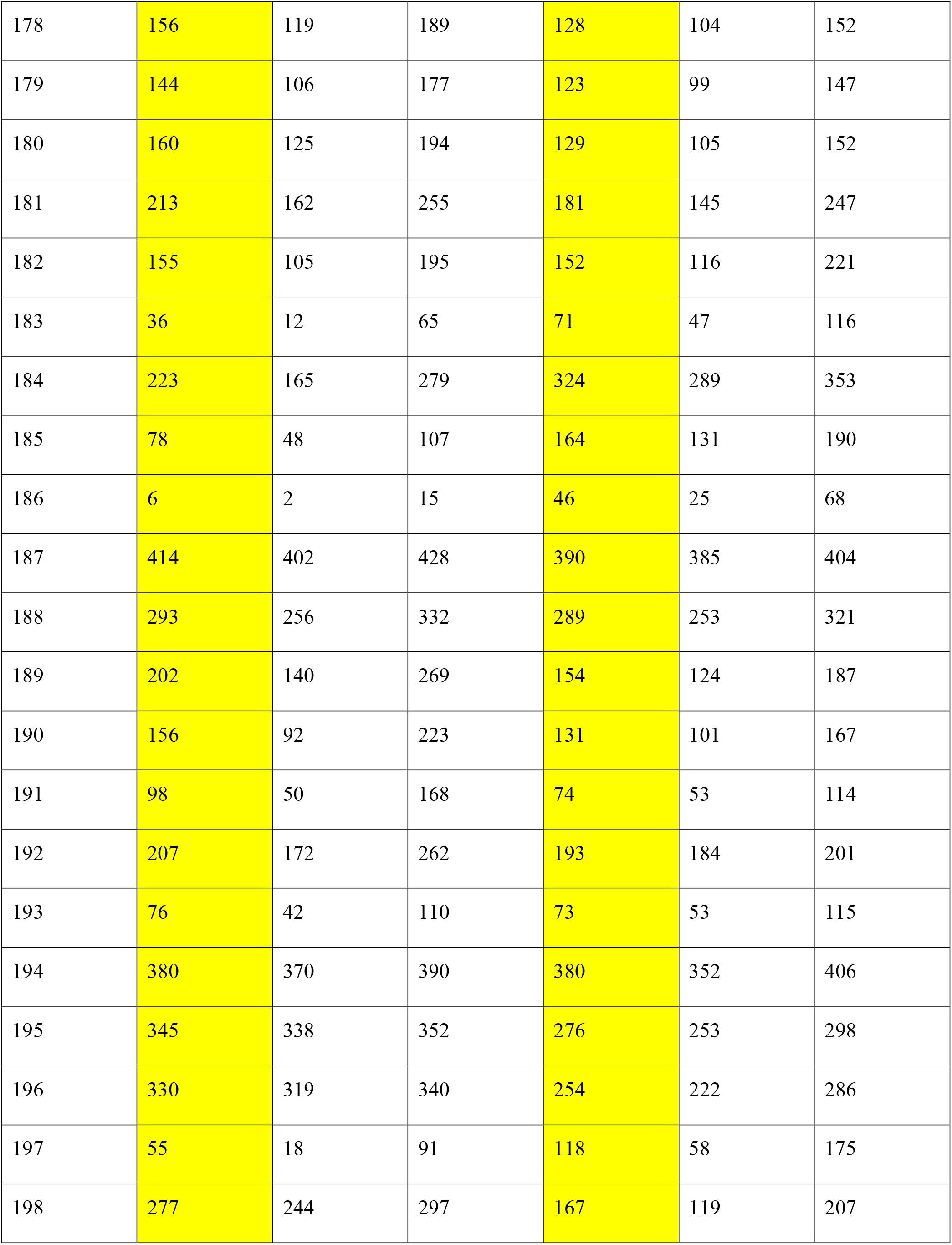
The ages found in PhyloBayes by Irisarri et al. (2017: supplementary table 9: last three columns) when all calibrations were used (all bounds treated as soft, mean ages averaged over 100 gene-jackknifed runs, extremes absolute over all runs), and the results obtained here in PhyloBayes with the updated calibrations (all bounds treated as hard, mean ages averaged over two runs with the full dataset, extremes absolute over both runs). All calibration dates are shown in Table 1. All ages are rounded to whole Ma. CI = credibility interval.

| Node number | Irisarri et al.<br>(2017) |  |  | Present<br>results |  |  |
| --- | --- | --- | --- | --- | --- | --- |
|  | Mean age | younger end<br>of 95% CI | older end of<br>95% CI | Mean age | younger end<br>of 95% CI | older end of<br>95% CI |
| 100 | 460 | 452 | 465 | 472 | 467 | 475 |
| 101 | 393 | 383 | 403 | 390 | 363 | 415 |
| 102 | 437 | 431 | 440 | 462 | 453 | 471 |
| 103 | 426 | 420 | 431 | 435 | 427 | 445 |
| 104 | 412 | 408 | 418 | 423 | 420 | 4230 |
| 105 | 341 | 331 | 350 | 363 | 359 | 365 |
| 106 | 289 | 283 | 296 | 320 | 318 | 324 |
| 107 | 257 | 256 | 257 | 301 | 294 | 307 |
| 108 | 254 | 253 | 256 | 290 | 282 | 298 |
| 109 | 243 | 242 | 245 | 250 | 248 | 252 |
| 110 | 120 | 90 | 162 | 162 | 129 | 185 |
| 111 | 71 | 66 | 75 | 160 | 126 | 182 |
| 112 | 137 | 111 | 173 | 167 | 141 | 186 |
| 113 | 83 | 70 | 87 | 105 | 88 | 115 |
| 114 | 63 | 47 | 73 | 90 | 70 | 102 |
| How to calibrate gnathostome timetrees |  |  |  |  |  |  |
| 115 | 16 | 8 | 25 | 66 | 52 | 80 |
| 116 | 92 | 66 | 130 | 163 | 136 | 183 |
| 117 | 224 | 211 | 234 | 181 | 175 | 185 |
| 118 | 206 | 184 | 221 | 167 | 148 | 178 |
| 119 | 168 | 133 | 188 | 142 | 112 | 157 |
| 120 | 155 | 117 | 176 | 140 | 109 | 155 |
| 121 | 127 | 90 | 150 | 135 | 104 | 151 |
| 122 | 95 | 63 | 124 | 116 | 86 | 136 |
| 123 | 78 | 45 | 107 | 120 | 91 | 139 |
| 124 | 192 | 167 | 211 | 172 | 160 | 181 |
| 125 | 239 | 233 | 244 | 254 | 244 | 268 |
| 126 | 199 | 190 | 208 | 153 | 141 | 178 |
| 127 | 195 | 185 | 204 | 150 | 138 | 175 |
| 128 | 187 | 177 | 196 | 144 | 133 | 170 |
| 129 | 182 | 173 | 192 | 141 | 131 | 167 |
| 130 | 181 | 172 | 190 | 139 | 128 | 165 |
| 131 | 166 | 159 | 175 | 131 | 119 | 157 |
| 132 | 137 | 124 | 151 | 117 | 101 | 142 |
| 133 | 127 | 111 | 142 | 115 | 99 | 141 |
| 134 | 130 | 115 | 145 | 92 | 67 | 117 |
| 135 | 128 | 104 | 143 | 113 | 95 | 138 |
| 136 | 94 | 72 | 119 | 105 | 91 | 131 |
| 137 | 88 | 66 | 112 | 102 | 86 | 128 |
| 138 | 64 | 40 | 91 | 82 | 69 | 106 |
| 139 | 47 | 26 | 72 | 72 | 59 | 93 |
| 140 | 11 | 4 | 25 | 60 | 46 | 79 |
| 141 | 46 | 25 | 72 | 77 | 64 | 100 |
| 142 | 27 | 13 | 49 | 52 | 37 | 69 |
| 143 | 39 | 21 | 64 | 74 | 61 | 97 |
| 144 | 22 | 11 | 42 | 72 | 58 | 94 |
| 145 | 179 | 167 | 190 | 106 | 82 | 134 |
| 146 | 156 | 136 | 172 | 116 | 103 | 143 |
| 147 | 57 | 34 | 77 | 69 | 59 | 93 |
| 148 | 44 | 24 | 65 | 69 | 58 | 93 |
| 149 | 165 | 146 | 181 | 138 | 123 | 164 |
| 150 | 165 | 161 | 172 | 229 | 217 | 233 |
| 151 | 138 | 136 | 140 | 142 | 128 | 157 |
| 152 | 94 | 91 | 96 | 71 | 69 | 72 |
| 153 | 89 | 85 | 92 | 68 | 67 | 70 |
| 154 | 61 | 53 | 65 | 54 | 50 | 56 |
| 155 | 79 | 71 | 84 | 66 | 65 | 67 |
| 156 | 91 | 87 | 94 | 64 | 56 | 69 |
| How to calibrate gnathostome timetrees |  |  |  |  |  |  |
| 157 | 68 | 62 | 72 | 66 | 61 | 68 |
| 158 | 50 | 38 | 60 | 59 | 51 | 67 |
| 159 | 315 | 300 | 328 | 328 | 314 | 339 |
| 160 | 307 | 290 | 323 | 286 | 275 | 290 |
| 161 | 202 | 173 | 237 | 170 | 137 | 188 |
| 162 | 192 | 163 | 226 | 165 | 131 | 183 |
| 163 | 177 | 146 | 210 | 143 | 110 | 160 |
| 164 | 168 | 137 | 199 | 139 | 106 | 156 |
| 165 | 117 | 86 | 143 | 104 | 71 | 119 |
| 166 | 92 | 62 | 117 | 60 | 36 | 70 |
| 167 | 77 | 49 | 101 | 59 | 35 | 69 |
| 168 | 53 | 30 | 74 | 45 | 26 | 56 |
| 169 | 162 | 134 | 196 | 143 | 107 | 163 |
| 170 | 201 | 170 | 232 | 173 | 156 | 193 |
| 171 | 192 | 161 | 224 | 170 | 154 | 192 |
| 172 | 186 | 154 | 218 | 166 | 148 | 188 |
| 173 | 155 | 123 | 186 | 147 | 127 | 172 |
| 174 | 105 | 71 | 140 | 104 | 85 | 141 |
| 175 | 94 | 62 | 127 | 74 | 57 | 109 |
| 176 | 70 | 33 | 110 | 75 | 60 | 117 |
| 177 | 54 | 22 | 89 | 71 | 56 | 111 |
| 178 | 156 | 119 | 189 | 128 | 104 | 152 |
| 179 | 144 | 106 | 177 | 123 | 99 | 147 |
| 180 | 160 | 125 | 194 | 129 | 105 | 152 |
| 181 | 213 | 162 | 255 | 181 | 145 | 247 |
| 182 | 155 | 105 | 195 | 152 | 116 | 221 |
| 183 | 36 | 12 | 65 | 71 | 47 | 116 |
| 184 | 223 | 165 | 279 | 324 | 289 | 353 |
| 185 | 78 | 48 | 107 | 164 | 131 | 190 |
| 186 | 6 | 2 | 15 | 46 | 25 | 68 |
| 187 | 414 | 402 | 428 | 390 | 385 | 404 |
| 188 | 293 | 256 | 332 | 289 | 253 | 321 |
| 189 | 202 | 140 | 269 | 154 | 124 | 187 |
| 190 | 156 | 92 | 223 | 131 | 101 | 167 |
| 191 | 98 | 50 | 168 | 74 | 53 | 114 |
| 192 | 207 | 172 | 262 | 193 | 184 | 201 |
| 193 | 76 | 42 | 110 | 73 | 53 | 115 |
| 194 | 380 | 370 | 390 | 380 | 352 | 406 |
| 195 | 345 | 338 | 352 | 276 | 253 | 298 |
| 196 | 330 | 319 | 340 | 254 | 222 | 286 |
| 197 | 55 | 18 | 91 | 118 | 58 | 175 |
| 198 | 277 | 244 | 297 | 167 | 119 | 207 |

### 2.2 Calibrations

In the 29 subsections below and in the Supplementary Material I discuss the minimum and maximum ages of all 30 nodes used as calibrations by Irisarri et al. (2017), referring to each by clade names and by the node number assigned by Irisarri et al. (2017: especially supp. table 8 and supp. fig. 19), also shown in Fig. 1. The abbreviation Fm stands for Formation; ICSC refers to the International Chronostratigraphic Chart v2020/3 (Cohen et al., 2020a); Ma is the quasi-SI symbol for megayear (million years).

#### 2.2.1 Root node (100): Gnathostomata [PN] (total group including Chondrichthyes – Pan-Osteichthyes [PN])

The cladogenesis that created the total groups of Chondrichthyes and Osteichthyes [PN] was assigned a minimum age of 421.75 Ma, a remarkably precise date close to the Silurian-Devonian boundary, and a maximum age of 462.5 Ma, which is currently (ICSC) thought to lie in the Darriwilian stage of the Middle Ordovician.

The Darriwilian should rather be regarded as the minimum age of this calibration date. While articulated bones and teeth of gnathostomes – both total-group chondrichthyans (Burrow and Young, 1999) and pan-osteichthyans (Choo et al., 2017, and references therein) – are only known from the Ludfordian (Ludlow, late Silurian) upward, a large diversity of scales that are increasingly confidently assigned to stem-chondrichthyans extends all the way down into the early Darriwilian (Sansom et al., 2012; Andreev et al., 2015, 2016a, b; Žigaitė Andreev, 2018; and references therein). The Darriwilian is currently thought to have begun 467.3 ± 1.1 Ma ago and to have ended 458.4 ± 0.9 Ma ago (ICSC); for the purposes of reducing “the middle part of the Stairway Sandstone” (Sansom et al., 2012: 243) to a single number, the age of 465 Ma should be adequate as the minimum age of Gnathostomata.

As a maximum age I cautiously propose the mid-Floian (Early Ordovician) upper fossiliferous level of the Burgess-like Fezouata Shale; at both levels, gnathostomes are absent among the “over 200 taxa, about half of which are soft-bodied” (Lefebvre et al., 2017: 296). Note that the oldest known hard tissues of vertebrates are Floian in age as well (reviewed by Sansom and Andreev, 2018). The Floian began 477.7 ± 1.4 Ma ago and ended 470.0 ± 1.4 Ma ago (ICSC), so I suggest a soft maximum age of 475 Ma for this calibration date.

The minimum and the maximum age proposed here are unexpectedly close together. This may be a sign that one or both is an unduly optimistic assessment of our knowledge of the fossil record – or that the origin of Gnathostomata formed part of the Great Ordovician Biodiversification Event (Sansom et al., 2012; Sansom and Andreev, 2018), which does not seem implausible.

#### 2.2.2 Node 102: Osteichthyes [PN] (Pan-Actinopterygii [PN] – Sarcopterygii)

Irisarri et al. (2017) assigned a minimum age of 416 Ma and a maximum age of 439 Ma, spanning the Silurian-Devonian boundary, to the cladogenesis that created the osteichthyan crown-group by separating the sister-groups Pan-Actinopterygii and Sarcopterygii.

The minimum age of this cladogenesis event depends on the phylogenetic position of the “psarolepids” (Choo et al., 2017) *Guiyu* and *Sparalepis* from the Kuanti [Guandi] Fm of Yunnan, China, which represents an early part of the abovementioned Ludfordian stage (425.6 ± 0.9 to 423.0 ± 2.3 Ma ago: ICSC). The “psarolepids” lie either just outside Osteichthyes or just inside, on the sarcopterygian side of the basal dichotomy (Clement et al., 2018, and references therein). To some extent the result depends on the analysis method: Clement et al. (2018) found the “psarolepids” outside Osteichthyes by parsimony (bootstrap support throughout the tree artificially low due to missing data), but inside by Bayesian inference (94% posterior probability). Following the discussions of this issue in Choo et al. (2017), Lu et al. (2017) and Clement et al. (2018), and in particular the work of King (2019), I favor a stem-pan-osteichthyan position for this assemblage over a large number of unexpected reversals to a “placoderm” state.

The oldest known uncontroversial osteichthyan is the oldest known dipnomorph, *Youngolepis*, as discussed below; following the assignment of *Andreolepis* and *Lophosteus* to the osteichthyan stem (e.g. Botella et al., 2007; Chen et al., 2016), all certain or uncertain actinopterygians are Devonian or younger. Thus, the minimum age for this calibration is the same as that for the next, Node 104.

Likewise, for the same reasons as discussed under Node 104, I cannot assign a maximum age to this divergence other than that for the root node. I have, in other words, not calibrated this node, and recommend against using this cladogenetic event as a calibration date if Nodes 100 and 104 are available.

#### 2.2.3 Node 104: Dipnomorpha – Tetrapodomorpha

The divergence of the sister-groups Dipnomorpha (the lungfish total group) and Tetrapodomorpha (the tetrapod total group) was assigned a minimum age of 408 and a maximum age of 419 Ma.

The minimum age may not contradict the age of the oldest known tetrapodomorph, *Tungsenia*, which is Pragian in age (Lu et al., 2012); the beginning of the Pragian is dated to 410.8 ± 2.8 Ma, its end to 407.6 ± 2.6 Ma (ICSC). However, the minimum age is clearly younger than the oldest known dipnomorphs. The oldest known specimens have been referred to *Youngolepis* and come from the lower part of the Xishancun Fm (Zhu and Fan, 1995). This formation is generally (e.g. Choo et al., 2017; Liu et al., 2017; and references therein) considered to represent the lower third or less of the Lochkovian stage, its bottom coinciding with the Silurian-Devonian boundary, which is currently dated to 419.2 ± 3.2 Ma (ICSC). However, Zhang et al. (2014) placed it in the middle of the immediately preceding Přídolí stage, which began 423.0 ± 2.3 Ma ago (ICSC). Needing a single number to summarize this uncertainty, I suggest a minimum age of 420 Ma for Node 104, the divergence of Dipnomorpha and Tetrapodomorpha. (This is a revision stratigraphically downward from the 410 Ma recommended by Marjanović and Laurin, 2007.)

A maximum age is difficult to assign. The abovementioned Kuanti Fm, which is universally (Zhang et al., 2014) regarded as representing an early part of the Ludfordian stage which preceded the Přídolí, has yielded several gnathostomes, but the sample seems too small to tell whether the absence of dipno- and tetrapodomorphs is real. Only one even partial articulated gnathostome is known from any other Ludfordian site in the world (*Yealepis*, which lies on the chondrichthyan stem: Burrow and Young, 1999). Comparably rich sites older than the Ludfordian have not been discovered. I cannot recommend any particular maximum age for this calibration point, other than by implication the maximum age of the root node (475 Ma, see above).

#### 2.2.4 Node 105: Tetrapoda [PN] (Amphibia [PN] – Pan-Amniota [PN])

The divergence between the ancestors of lissamphibians and those of amniotes was assigned a minimum age of 330.4 and a maximum of 350.1 Ma following Benton and Donoghue (2007). Although Pardo et al. (2020) have reviewed the breadth of issues it raises far beyond the scope of this work, and I broadly agree with their conclusions, a few points still remain to address or summarize.

For a long time, the oldest tetrapod was thought to be *Lethiscus*, variably supposed to be a stem-amphibian or a stem-pan-amniote (see below), which is mid-Viséan in age (Smithson et al., 2012, and references therein; the Viséan lasted from 346.7 ± 0.4 to 330.9 ± 0.2 Ma ago: ICSC). More likely, *Lethiscus* and the other aïstopods are rather early-branching stem-stegocephalians [PN] (Pardo et al., 2017, 2018; Clack et al., 2019; further discussion in Marjanović and Laurin, 2019). Whether *Casineria* from a geographically (southeastern Scotland) and stratigraphically close site (mid-late Viséan: Paton et al., 1999; Smithson et al., 2012) can replace it in that function depends on two unresolved issues: its own phylogenetic position, for which estimates range from very close to Amniota (within Tetrapoda) into Temnospondyli (Marjanović and Laurin, 2019, and references therein; Clack et al., 2019; Daza et al., 2020: fig. S15); and the controversial phylogenetic position of Lissamphibia [PN] in the stegocephalian tree (Marjanović and Laurin, 2013a, 2019; Danto et al., 2019; Laurin et al., 2019; Pardo et al., 2020; Daza et al., 2020; and references in all five), which determines whether the temnospondyls are tetrapods or quite rootward stem-stegocephalians by determining which node of the otherwise largely stable tree of early stegocephalians bears the name Tetrapoda.

Anderson et al. (2015) reported a number of isolated anthracosaur [PN] (embolomere or eoherpetid) bones from a mid-Tournaisian site (the Tournaisian preceded the Viséan and began at the Devonian/Carboniferous boundary 358.9 ± 0.4 Ma ago: ICSC). Whether these are tetrapods depends on the relative positions of temnospondyls, anthracosaurs and other clades in that region of the tree (Pardo et al., 2018, 2020; Marjanović and Laurin, 2019; Ruta et al., 2020; and references in all four) in addition to the position of Lissamphibia: even if the lissamphibians are temnospondyls, the anthracosaurs may still be stem-stegocephalians.

The same site has also yielded the oldest colosteid remains (Anderson et al., 2015). Colosteidae (“Colosteida” of Pardo et al., 2020) was referred to Temnospondyli throughout the 20^th^ century and found in that position by Marjanović trees by Daza et al., 2020: fig. S15) and Laurin (2019) to our great surprise (also in some of the treess pointed out by Pardo et al. (2020), this means it could belong to Tetrapoda. However, ongoing work on enlarging and improving the matrix of Marjanović and Laurin (2019) and Daza et al. (2020) shows this result was most likely an artefact of the taxon and character sample; similarly, Ruta et al. (2020) found the colosteid they included to be a temnospondyl with weak support in their Bayesian analysis, but to lie rootward of Temnospondyli in their parsimony analyses (unweighted, reweighted or with implied weighting).

The same site has further yielded tetrapod trackways, some of which are tetradactyl (Smithson et al., 2012, and references therein). Among Paleozoic tetrapods, tetradactyly is only known among “microsaurs” (including lysorophians), scincosaurids, some urocordylids, temnospondyls and *Colosteus* (but not its close pentadactyl relative *Greererpeton*). (Reports of tetradactyl limbs in diplocaulids have been erroneous: Milner, 2019; Marjanović and Laurin, 2019, and references therein.) *Colosteus* and probably (Clack et al., 2019) the urocordylids are stem-stegocephalians, but both were fully aquatic, thus unlikely to leave trackways; “microsaurs” and probably scincosaurids were tetrapods, and most were amphibious to terrestrial; temnospondyls spanned the full range of lifestyles, but see above for their phylogenetic position. In short, whether tetradactyl trackways are evidence of tetrapods in the mid-late Tournaisian remains unclear.

The oldest uncontroversial tetrapod is thus *Westlothiana* from close to the end of the Viséan (Marjanović and Laurin, 2019, and references therein, especially Smithson et al., 1994, 2012). Other stegocephaans from the same site and age may or may not be tetrapods: whether the temnospondyl *Balanerpeton* (Milner and Sequeira, 1994; Schoch and Milner, 2014) is one depends on the resolution of the abovementioned controversy about Lissamphibia; likewise, see above on the “anthracosaur-grade” (Marjanović and Laurin, 2019; Ruta et al., 2020) animals *Silvanerpeton* and *Eldeceeon*; *Ophiderpeton kirktonense* is an aïstopod, on which see above; *Kirktonecta* (Clack, 2011) is likely a tetrapod, but needs to be fully prepared or μ be made.

Thus, the minimum age may be as young as roughly 335 Ma (mid-late Viséan) or as old as roughly 350 Ma (early-middle Tournaisian) depending on two phylogenetic problems.

The few Tournaisian tetrapod sites discovered so far (Smithson et al., 2012; Anderson et al., 2015; Clack et al., 2016) have not yielded any uncontroversial tetrapods, temnospondyl bones or temnospondyl footprints; thus, if the temnospondyls are stem-tetrapodomorphs, the ages of these sites (up to roughly 350 Ma) may be useful as a maximum age. However, as stressed by Pardo et al. (2020), they represent a very small region of the Carboniferous globe, so I continue (Marjanović and Laurin, 2019) to caution against this regardless of the phylogenetic issues. Rather, the richer and better studied Famennian (end-Devonian) record, which has not so far yielded close relatives of Tetrapoda but has yielded more rootward stegocephalians and other tetrapodomorphs (Marjanović and Laurin, 2019; Ahlberg and Clack, 2020; and references therein), should be used to place a soft maximum age around very roughly 365 Ma.

#### 2.2.5 Node 106: Amniota [PN] (Pan-Mammalia [PN] – Sauropsida)

The cladogenesis that separated the total group of mammals (also called Synapsida [PN] or Theropsida: Goodrich, 1916) from the total group of diapsids including turtles (Sauropsida: Goodrich, 1916) was assigned a minimum age of 288 Ma (Artinskian, Early Permian) and a maximum age of 338 Ma (Viséan, Early Carboniferous).

This minimum age is rather puzzling. I am not aware of any doubts on the membership of *Hylonomus* in Sauropsida since its redescription by Carroll (1964), except the very vague ones presented by Graur and Martin (2004) and taken from even more outdated literature; none are mentioned in the review by Pardo et al. (2020) either. Because of its late Bashkirian age, this calibration has often been dated to 310 Ma (as discussed by Graur and Martin, 2004). Currently (ICSC), the Bashkirian is thought to have ended 315.2 ± 0.2 and begun 323.2 ± 0.4 Ma ago, and the site (Joggins, Nova Scotia) that has yielded *Hylonomus* has been dated to 317–319 Ma (Carpenter et al., 2015); thus, given the phylogenetic position of *Hylonomus* (Ford and Benson, 2019, and references therein), I suggest a minimum age of 318 Ma for this calibration.

There appears to be pan-mammalian material from the same site (Carroll, 1964; Mann et al., 2020), which has also yielded various “microsaurs” that Pardo et al. (2017) included in Sauropsida (see also Marjanović and Laurin, 2019, and Pardo et al., 2020). I should also emphasize that the next younger sauropsids and pan-mammals (and “microsaurs”) older than 288 Ma come from several sites in each following geological stage (Moscovian through Artinskian) and represent a considerable diversity; from the Moscovian alone, four sites of successive ages are known that present more or less complete skeletons of uncontroversial amniotes, namely sauropsids closely related to Diapsida and *Hylonomus* (*Anthracodromeus*, *Brouffia*, *Cephalerpeton*, *Paleothyris*), the oldest “parareptile” (*Carbonodraco*) as well as what appears to be the sister-group to most other sauropsids (*Coelostegus*), and, on the pan-mammalian side, ophiacodontids (*Echinerpeton*; *Archaeothyris* from two sites). A fifth site preserves the oldest varanopid, a group of amniotes of unclear phylogenetic position (Ford and Benson, 2018, 2019). As reviewed in detail by Pardo et al. (2020), this implies ghost lineages for several other amniote clades that might not have lived in coal swamps; several of these show up in the fossil record of the next and last two stages of the Carboniferous, which ended 298.9 ± 0.15 Ma ago (ICSC). For more information on the Carboniferous amniote record see Reisz and Modesto (1996: fig. 3), Müller and Reisz (2006), Mann and Paterson (2019), Mann et al. (2019), Maddin et al. (2019) and Pardo et al. (2020), the second and the third with phylogenetic analyses, as well as references in all six. Additionally, the oldest known diadectomorphs (“diadectamorphs” of Pardo et al., 2020) date from the Kasimovian (“Missourian” in Kissel, 2010) which follows the Moscovian; they may represent the sister-group of Amniota, or they may be what should have been called non-synapsid theropsids (Marjanović and Laurin, 2019; Klembara et al., 2019; Pardo et al., 2020; and references in all three).

The absence of amniotes (and diadectomorphs) in the Serpukhovian record preceding the Bashkirian should not be given much weight for paleoecological reasons, as reviewed by Pardo et al. (2020); note that “lepospondyls” like the Viséan *Kirktonecta* and *Westlothiana*, probably closely related to but outside Amniota, are almost unknown from this age as well (candidates were described by Carroll et al., 1991; Carroll and Chorn, 1995; Lombard and Bolt, 1999). Their absence from the somewhat richer Viséan record (discussed above) suffers in part from the same problem, in part from geographic restrictions. Thus, I refrain from recommending a maximum age other than that of the preceding Node 105, even though such an early age would imply very slow rates of morphological evolution in the earliest pan-mammals and sauropsids.

#### 2.2.6 Node 107: Reptilia [PN] (Pan-Lepidosauria – total group of Archelosauria); node 108: Archelosauria (Pan-Testudines [PN] – Pan-Archosauria [PN])

The origin of the sauropsid crown group by a split into Pan-Lepidosauria and the total group of Archelosauria was assigned a minimum age of 252 Ma and a maximum age of 257 Ma, both in the Late Permian. Ezcurra et al. (2014; correction: The PLOS ONE Staff, 2014) agreed that the oldest unambiguous reptile that can be clearly dated is the supposed pan-archosaur *Protorosaurus*, which is, however, 257.3 ± 1.6 Ma old as they also discussed. Therefore, they revised the minimum age to255.7 Ma, the younger end of this confidence interval.

However, like all other recent phylogenetic analyses of molecular data, Irisarri et al. (2017) found the turtles to be closer to Archosauria [PN] than Lepidosauria [PN]. Thus, the question whether *Eunotosaurus* is a member of the turtle stem (Schoch and Sues, 2017, and references therein) becomes relevant, because the earliest occurrence of *Eunotosaurus* is roughly middle Capitanian in age (the Capitanian, the last stage of the Middle Permian, ended 259.1 ± 0.5 Ma ago and began 265.1 ± 0.4 Ma ago: ICSC), and further because *Protorosaurus* would presumably belong to Pan-Archosauria and thus calibrate Node 108, not 107.

For present purposes I set the minimum age of Archelosauria (Node 108) as 263 Ma, the approximate midpoint of the Capitanian, and do not assign a minimum age to Reptilia (Node 107). But in general I have to, at our current level of understanding, recommend against using either of these nodes as a calibration. The reason are two major uncertainties about the topology of the phylogenetic tree.

First, if *Eunotosaurus* has moved from the “parareptiles” well outside Diapsida [PN] – or well inside Diapsida, though presumably still in its stem-group (Ford and Benson, 2019) – to the turtle stem within the crown group of Diapsida (i.e. Reptilia [PN]), do any other “parareptiles” follow it? The oldest known member of that assemblage, *Carbonodraco*, comes from the site of Linton in Ohio (Mann et al., 2019), which is about 307–308 Ma old (compare Reisz and Modesto, 1996, and Carpenter et al., 2015), so that should be the minimum age of Archelosauria if all “parareptiles” are archelosaurs; the currently available phylogenetic analyses of “parareptiles” (Laurin and Piñeiro, 2018; MacDougall et al., 2019) have not adequately tested this question. While Schoch and Sues (2017) did test the mutual relationships of “parareptiles”, *Eunotosaurus* and diapsids and found *Eunotosaurus* nested in the latter, several nodes away from the former, these nodes were very poorly supported. The character and taxon samples of all existing matrices for analyses of amniote phylogeny need to be substantially improved (Ford and Benson, 2018, 2019; Laurin and Piñeiro, 2018; MacDougall et al., 2019; Mann et al., 2019); Ford and Benson (2019) made a large step in that direction, but deliberately excluded *Eunotosaurus* and the turtles from their analysis so as not to have to deal with all problems at the same time.

Second, the position of *Protorosaurus* as a pan-archosaur, accepted for decades, was thrown into doubt by Simões et al. (2018), who found it as such in their Bayesian analyses of morphological or combined data (Simões et al., 2018: ext. data fig. 5, 6; also, after a few changes to the dataset, Garberoglio et al., 2019: fig. S2; Sobral et al., 2020: fig. S9, S10), but not in their parsimony analyses of morphological data without or with implied weights (ext. data fig. 3, 4; likewise Garberoglio et al,. 2019: fig. S3, and Sobral et al., 2020: fig. S7, S8), where it came out as a stem-sauropsid; the question was unresolved in their Bayesian tip-dating or tip-and-node dating analyses of combined data (ext. data fig. 7, 8). After a different set of changes to the dataset, Simões et al. (2020) found *Protorosaurus* as a pan-archosaur when they used MrBayes (supp. fig. 2–5) or when they used BEAST for dating with a correction (supp. fig. 7), but not when they used BEAST for dating without a correction (supp. fig. 6). Support was moderate throughout. However, these trees are hard to compare to that of Irisarri et al. (2017) because they all find the turtles outside the diapsid crown (with limited support); no extant archosaurs or turtles, and therefore no molecular data for them, are included in these datasets. Using a smaller dataset with much denser sampling of Triassic reptiles, Pritchard et al. (2018) found *Protorosaurus* closer to Archosauria than to Lepidosauria with very strong support (parsimony bootstrap value: 100%, Bayesian posterior probability: 99.06%), but whether that is on the archosaur or the archelosaur stem could not be determined because there were no turtles in that dataset.

The maximum age of either node is likewise difficult to narrow down. Uncontroversial diapsids have a notoriously patchy Paleozoic record (Ford and Benson, 2018, and references therein); the same holds for “parareptiles”, which have only two known Carboniferous records so far (Modesto et al., 2015; Mann et al., 2019). I cannot express confidence in a maximum age other than that of Node 106, which I cannot distinguish from the maximum age of Node 105 as explained above. This leaves Node 107 without independent calibrations in the current taxon sample.

#### 2.2.7 Node 109: Archosauria [PN] (Crocodylotarsi – Pan-Aves [PN])

The origin of Archosauria by cladogenesis into the total groups of crocodiles and birds was given a minimum age of 243 Ma (Middle Triassic) and a maximum age of 251 Ma (Early Triassic).

The earliest securely dated known archosaur, belonging to the crocodile stem, is *Ctenosauriscus* from just before the end of the Olenëkian; several close relatives may be coeval or a little younger (Butler et al., 2011). The age of the Olenëkian/Anisian (Early/Middle Triassic) boundary is given in the ICSC as 247.2 Ma without a confidence interval; any such confidence interval cannot be long, however, because an Olenëkian sample has been dated to 247.32 ± 0.08 Ma, while an Anisian sample has been dated to 247.08 ± 0.11 Ma (Maron et al., 2018). Given the highly nested phylogenetic position of *Ctenosauriscus* in Archosauria (Butler et al., 2011; Ezcurra et al., 2020: ext. data fig. 4, 8), I propose 248 Ma as the minimum age of this calibration.

I accept the Permian-Triassic boundary (251.902 ± 0.024 Ma: ICSC; rounded to 252) as the soft maximum age on the grounds that a major radiation of pan-archosaurs at the beginning of the Triassic seems likely for ecological reasons: the Permian record, up to its very end, is full of pan-mammals that seem ecologically comparable to Triassic archosaurs, and given the Pangea situation of the time it seems reasonably unlikely that archosaurs existed in unsampled localities. I must caution, however, that the fossil record of pan-archosaurs and possible pan-archosaurs in the four million years of the Triassic preceding the minimum age, and in the Permian, is very patchy, with a poor fit between stratigraphy and phylogeny; indeed, the Permian record of archosauriforms [PN] is currently entirely limited to the poorly known non-archosaur *Archosaurus* and possibly the even more poorly known non-archosaur *Eorasaurus* (Ezcurra et al., 2014).

#### 2.2.8 Node 111: Alligatoridae (Alligatorinae – Caimaninae)

The origin of Alligatoridae (the crown group of Globidonta) by split into Alligatorinae and Caimaninae was given a minimum age of 66 Ma (the Cretaceous/Paleogene boundary) and a maximum age of 75 Ma (Campanian, Late Cretaceous).

The minimum age would fit well with the finding by Cossette and Brochu (2018) that *Bottosaurus* from the very end of the Cretaceous is a caimanine. Given, however, the limited material and the stratigraphic gap between *Bottosaurus* and the next younger known caimanines, Cossette and Brochu (2018) expressed doubt about the result of their phylogenetic analysis which placed *Bottosaurus* not only within the caimanine crown-group but next to the extant *Paleosuchus*. Cossette and Brochu (2020) did not include *Bottosaurus* in their phylogenetic analysis.

If *Bottosaurus* is not an alligatorid at all, the oldest known member is the alligatorine *Navajosuchus* from within the first million years of the Paleocene (Puercan NALMA [North American Land Mammal Age]), translating to a minimum age of 65 Ma (Wang et al., 2016, and references therein). The oldest known caimanines (*Protocaiman*, *Eocaiman paleocenicus* and *Necrosuchus*: Bona et al., 2018) follow shortly thereafter (Peligran SALMA [South American Land Mammal Age], 64–63 Ma ago: Woodburne et al., 2014).

Halliday et al. (2013), however, found the Campanian to Maastrichtian *Brachychampsa* to be an alligatorine, as did Arribas et al. (2019) in a less densely sampled analysis of Crocodyliformes; Bona et al. (2018) found it and the newly added Campanian *Albertochampsa* to be caimanines, a finding expanded by Cossette and Brochu (2020) to *Stangerochampsa*. In all these cases, the earliest record of an alligatorid is *Brachychampsa sealeyi* from early in the Campanian, which began 83.6 ± 0.2 Ma ago (ICSC). These results were not replicated by Lee and Yates (2018) or by Groh et al. (2019), who both found *Brachychampsa* on the brevirostrine stem, not as an alligatorid, and who both did not include *Albertochampsa* in their datasets. I must caution, however, that Groh et al. (2019) found Alligatorinae, and even *Alligator* itself, as a Hennigian comb in which Caimaninae was nested; this result strongly suggests that the character sample was insufficient to resolve Brevirostres.

Given this uncertainty, I have used a minimum age of 65 Ma for present purposes, but generally recommend against using this cladogenesis as a calibration for timetrees.

Up to (and including) the Campanian, the record of neosuchians is a surprisingly spotty affair (e.g. Tykoski et al., 2002; Mateus et al., 2018). Although a Late Cretaceous age of Alligatoridae (i.e. less than 100.5 Ma: ICSC) is likely, I cannot, therefore, assign a maximum age younger than the Triassic/Jurassic boundary, i.e. twice as old (201.3 ± 0.2 Ma: ICSC; rounded to 200). Only in the Triassic is the record of ecologically comparable phytosaurs dense enough to really rule out the presence of amphibious crocodylomorphs such as alligatorids. However, I have treated this maximum as hard because the likelihood that the true age approaches it is very low.

#### 2.2.9 Node 113: Neognathae (Galloanserae [PN] – Neoaves)

The last common ancestor of *Anas*, *Gallus* and *Meleagris* on one side and *Taeniopygia* on the other was assigned a minimum age of 66 Ma (the Cretaceous/Paleogene boundary) and a maximum age of 86.5 Ma (Coniacian/Santonian boundary. Late Cretaceous) following Benton and Donoghue (2007).

The oldest known neognath appears to be the presbyornithid stem-anserimorph (Elżanowski, 2014; Tambussi et al., 2019; within two steps of the most parsimonious trees of Field et al., 2020) *Teviornis* from somewhere low in the Late Cretaceous Nemegt Fm of Mongolia; it is known only from a carpometacarpus, two phalanges and the distal end of a humerus that all seem to belong to the same right wing (Kurochkin et al., 2002). The most recent work on the specimen has bolstered its presbyornithid identity (De Pietri et al., 2016), even though the next younger presbyornithids are middle or late Paleocene (i.e. younger than 61.6 Ma: ICSC).

The age of the Nemegt Fm is difficult to pin down; radiometric dating of this or adjacent formations has not been possible, and the only fossils available for biostratigraphy are vertebrates that have to be compared to those of North America where marine correlations and radiometric dates are known. These comparisons favor a vaguely early Maastrichtian age, without ruling out a Campanian component. Magnetostratigraphic evidence was reported in a conference abstract by Hicks et al. (2001); I have not been able to find a follow-up publication. Hicks et al. (2001) stated that the sampled sections from the Nemegt and the conformably underlying Baruungoyot Fm “can be quite reliably correlated to the Geomagnetic Reversal Time Scale […] and clearly lie in the Campanian/Maastrichtian interval that extends from the uppermost part of subchron C33n, through chron 32 into the lower half of chron 31.” Where the Baruungoyot/Nemegt boundary lies on this scale was not mentioned. The upper boundary of the Nemegt Fm is an unconformity with a Paleocene formation.

Hicks et al. (2001) also studied the Late Cretaceous Djadokhta Fm, finding that “a distinct reversal sequence is emerging that allows us to correlate the sections in a preliminary way to the late Campanian through Maastrichtian interval that ranges from C32 to C31.” While I have not been able to find a publication by an overlapping set of authors on this finding, it agrees at least broadly with Dashzeveg et al. (2005: 18, 26, 27), whose own magnetostratigraphic work on the Djadokhta Fm indicated “that the sediments were deposited during the rapid sequence of polarity changes in the late part of the Campanian incorporating the end of Chron 33 and Chron 32 between about 75 and 71 Ma […]. However, this tentative correlation to the Geomagnetic Polarity Timescale cannot yet be certainly established.” Hasegawa et al. (2008) disagreed with the stratigraphy by Dashzeveg et al. (2005), but not with their dating.

Most often, the Djadokhta Fm has been thought to underlie the Baruungoyot Fm, but a contact between the two has not so far been identified (Dingus et al., 2008; cited without comment e.g. by Chinzorig et al., 2017); they could be partly coeval (references in Hasegawa et al., 2008). Still, it seems safe to say that most of the Nemegt Fm is younger than most of the Djadokhta Fm.

According to Milanese et al. (2018: fig. 12), the Campanian-Maastrichtian boundary (72.1 ± 0.2 Ma ago: ICSC) lies near the end of chron 32. The Djadokhta Fm thus corresponds to the end of the Campanian, the Baruungoyot Fm should have at most the same age, and the youngest magnetostratigraphic sample from the Nemegt Fm, in the earlier half of chron 31, should be about 70 Ma old. Given the stratigraphic position of *Teviornis* low within the formation and its nested phylogenetic position within Neognathae, I propose 71 Ma (within the same subchron as 70 Ma: Milanese et al., 2018: fig. 12) as the soft minimum age of the present calibration.

Field et al. (2020: 400) stated that the likely stem-pangallanseran “*Asteriornis* provides a firm calibration point for the minimum age of divergence of the major bird clades Galloanserae and Neoaves. We recommend that a minimum age of 66.7 million years is assigned to this pivotal neornithine node in future divergence time studies, reflecting the youngest possible age of the *Asteriornis* holotype including geochronological uncertainty.” In their supplementary information (p. 13), however, they revealed being aware of *Teviornis*, citing De Pietri et al. (2016) for its position as a presbyornithid (and thus, by their own phylogenetic analyses, an anserimorph) without discussing it any further.

Should the fragmentary *Teviornis* fall out elsewhere, the minimum age might nonetheless not have to rest on *Asteriornis*, because Vegaviidae, a clade containing the late Maastrichtian (Clarke et al., 2005; Salazar et al., 2010) *Vegavis*, *Polarornis* and *Neogaeornis* and probably the end-Campanian (McLachlan et al., 2017) *Maaqwi*, has been found on the anserimorph stem in some of the latest analyses (Agnolín et al., 2017; Tambussi et al., 2019). However, Mayr et al. (2018) discussed reasons for skepticism, and the analyses of McLachlan et al. (2017), Bailleul et al. (2019: supp. trees 7–11, 16, 17), Field et al. (2020) and O’Connor et al. (2020) found the vegaviids they included close to but outside Aves (or at least Galloanserae in the case of Bailleul et al., 2019, and O’Connor et al., 2020, who did not sample Neoaves or Palaeognathae in the analyses in question).

As the soft maximum age I tentatively suggest 115 Ma, an estimate of the mid-Aptian age of the terrestrial Xiagou Fm of northwestern China, which has yielded a diversity of stem-birds but no particularly close relatives of the crown (Wang et al., 2013; Bailleul et al., 2019; O’Connor et al., 2020; and references therein).

#### 2.2.10 Node 117: Testudines [PN] (Pan-Pleurodira [PN] – Pan-Cryptodira [PN])

The origin of the turtle crown group by split into the pleurodiran [PN] and cryptodiran [PN] total groups was assigned a minimum age of 210 Ma and no maximum age; this was taken from Noonan and Chippindale (2006) who cited a work from 1990 as their source.

The calibration dates treated above are almost all too young (some substantially so, others by just a few million years). This one, in contrast, is far too old. It rests on the outdated interpretation of the Norian (Late Triassic) *Proterochersis* as a stem-group pan-pleurodire. With one short series of exceptions (Gaffney et al., 2006, 2007; Gaffney and Jenkins, 2010), all 21^st^-century treatments of Mesozoic turtle phylogeny have found *Proterochersis* and all other turtles older than those mentioned below to lie well outside the crown group (Shao et al., 2018: fig. S8, S9; Sterli et al., 2019, 2020; and references therein, in Gaffney and Jenkins, 2010, and in Romano et al., 2014a).

The three oldest known xinjiangchelyids, of which one was referred to *Protoxinjiangchelys*, seem to be between 170 Ma and 180 Ma old (Aalenian/Bajocian boundary, Middle Jurassic, to Toarcian, late Early Jurassic; Hu et al., 2020, and reference therein). In the last three years, the xinjiangchelyids have been found as stem-testudinates or as stem-pan-cryptodires (Shao et al., 2018; Evers et al., 2019; González Ruiz et al., 2019: fig. 6, supp. fig. 4; Gentry et al., 2019; Anquetin and André, 2020; Sterli et al., 2020: supp. fig. “X” = 19), even in both positions when the same matrix was analyzed with different methods (Sterli et al., 2019: supp. file SterlietalSupplementary_material_3.pdf).

The oldest known securely dated and securely identified crown-group turtle is thus the mid-late Oxfordian stem-pan-pleurodire *Caribemys* (de la Fuente and Iturralde-Vinent, 2001; Shao et al., 2018; mostly referred to *Notoemys* as *N. oxfordiensis* in more recent literature, e.g. Sterli et al., 2019). Given that the Oxfordian ended 157.3 ± 1.0 Ma ago (ICSC), I suggest a minimum age of 158 Ma.

The stem-trionychian cryptodire *Sinaspideretes* (Tong et al., 2013), which would provide a minimum age for Cryptodira (node 118) rather than only Testudines, was long thought to have the same age or to be somewhat older. Of the three known specimens, at least one (the exact localities where the type and the other specimen were found are unknown) comes from the Upper (Shang-) Shaximiao Fm (Tong et al., 2013), which conformably overlies a sequence of two supposedly Middle Jurassic formations and is overlain by two Upper Jurassic formations (Tong et al., 2011; Xing et al., 2013), so it should be about Oxfordian to Callovian in age. The biostratigraphic evidence for the age of the Upper Shaximiao Fm is conflicting; there was no consensus on whether it is Middle or Late Jurassic (Xing et al., 2013) before Wang et al. (2018) showed that the immediately underlying Lower (Xia-) Shaximiao Fm is at most 159 ± 2 Ma old, a confidence interval that lies entirely in the Late Jurassic (which began, with the Oxfordian, 163.5 ± 1.0 Ma ago: ICSC). Most likely, then, the same holds for all *Sinaspideretes* specimens, and none of them is older than *Caribemys*.

The unambiguously Early Jurassic and Triassic record of turtles throughout Pangea lies entirely on the stem and has a rather good stratigraphic fit (see Sterli et al., 2019, 2020). I therefore suggest a soft maximum age of 185 Ma (in the Pliensbachian: ICSC) that probably postdates all of these taxa but predates the oldest possible age of the oldest known xinjiangchelyids.

#### 2.2.11 Node 124: Pleurodira [PN] (Pan-Chelidae – Pan-Pelomedusoides)

The origin of Pleurodira by the cladogenesis that generated Pan-Chelidae (represented by *Phrynops*) and Pan-Pelomedusoides (represented by *Pelusios*) was given a minimum age of 25 Ma (Oligocene) and no maximum age. This was miscopied from Noonan and Chippindale (2006: table 1), who assigned that age to Pelomedusidae (their calibration 18, represented here by *Pelusios* alone), not to Pleurodira; to Pleurodira they assigned (their calibration 17) a minimum age of 100 Ma (Early/Late Cretaceous boundary) and a maximum age of 150 Ma (Tithonian, Late Jurassic).

Pleurodira has long been known to extend into the Early Cretaceous (reviewed by Pérez-García, 2019); pan-podocnemidids within Pelomedusoides have a particularly rich fossil record. At present, the oldest known pleurodire is the late Barremian pan-podocnemidid *Atolchelys* (Romano et al., 2014a; Pérez-García, 2019; Hermanson et al., 2020), suggesting a minimum age of 125 Ma for this calibration (Romano et al., 2014a; ICSC).

Due to the fairly highly nested position of *Atolchelys* within Pleurodira (whether or not it is a bothremydid – Romano et al., 2014a; Cadena, 2015; Hermanson et al., 2020), and due to the somewhat sparse record of stem-pleurodires (from the Late Jurassic onwards: Romano et al., 2014a; Cadena, 2015; Pérez-García 2019), I accidentally agree with Irisarri et al. (2017) in not assigning a maximum age other than that of Node 117. The maximum age assigned by Noonan and Chippindale (2006: table 1) “assumes the Late Jurassic *Platychelys* actually predates the origin of modern pleurodiria [sic]”, which does not logically follow from the fact that it is close to but outside Pleurodira.

#### 2.2.12 Node 125: Lepidosauria [PN] (Rhynchocephalia – Pan-Squamata [PN])

The minimum age of this calibration, given as 238 Ma, has to be slightly revised to 244 Ma (both in the Middle Triassic) based on *Megachirella*, the oldest known unambiguous stem-pan-squamate (Renesto and Bernardi, 2013; Simões et al., 2018: table S2, 2020; Garberoglio et al., 2019; Sobral et al., 2020), which is older than the oldest known rhynchocephalian (238–240 Ma: Jones et al., 2013).

The Early Triassic *Sophineta*, a large collection of isolated bones, may be a stem-pan-squamate or a stem-pan-lepidosaur (Evans and Borsuk-Białynicka, 2009a; Simões et al., 2018, 2020; Garberoglio et al., 2019; Sobral et al., 2020). The text of Sobral et al. (2020) makes clear that *Vellbergia*, another such animal, is younger than *Megachirella*, despite being shown as older in Sobral et al. (2020: fig. 4).

An Early Triassic or perhaps Late Permian maximum age seems reasonable, but, given the rarity of stem-pan-lepidosaurs and of Permian diapsids in general (Carroll’s Gap – Marjanović and Laurin, 2013a), I rather propose to use the ecologically similar small amniotes (e.g. Haridy et al., 2017; MacDougall et al., 2019) of Richards Spur (up to 289.2 ± 0.68 Ma; Woodhead et al., 2010; MacDougall et al., 2017), immediately before Carroll’s Gap, to support a soft maximum age of 290 Ma.

#### 2.2.13 Node 129: Toxicofera (Pan-Serpentes [PN] – Anguimorpha + Pan-Iguania [PN])

This calibration was given a minimum age of 148 Ma (Tithonian, Late Jurassic) and no maximum age. Note that the minimum age was not operational because Node 131, Iguania, was given an older minimum age of 165 Ma; in other words, Node 129 was really not calibrated at all.

And indeed I should first mention that the pan-squamate fossil record suffers from three problems that make it difficult to calibrate this node. First, it exhibits Carroll’s Gap (Marjanović and Laurin, 2013a) very strongly. After the Middle Triassic stem-pan-squamate *Megachirella* and at least one Early Triassic pan-lepidosaur that may or may not be a pan-squamate (*Sophineta* in particular – compare the different phylogenetic analyses in Simões et al., 2018, 2020), the pan-squamate record as known today goes completely silent (see below under Node 131 for the one or two supposed exceptions) until the dam suddenly breaks in the Bathonian (Middle Jurassic) and representatives of the stem as well as, by current understanding, several parts of the crown appear in several sites in the northern continents and northernmost Gondwana. Second, these early representatives are all isolated and generally incomplete bones that preserve few diagnostic characters; the oldest complete skeletons come from one Tithonian (latest Jurassic) cluster of sites (Conrad, 2017), followed by a few Early Cretaceous ones as well as the oldest partially articulated material other than *Megachirella*. Third, the morphological datasets so far assembled for analysis of pan-squamate phylogeny are all so plagued by correlated characters and other problems that all of them support either Pan-Iguania as the sister-group to all other squamates, or the amphisbaenians (alone or even together with the dibamids) as the sister-group to Pan-Serpentes (e.g. Simões et al., 2020: supp. fig. 2), or both (e.g. Conrad, 2017: fig. 27, 28), while both are strongly contradicted by the molecular consensus (e.g. Irisarri et al., 2017; Garberoglio et al., 2019; Sobral et al., 2020: fig. S10; Simões et al., 2020: supp. fig. 1, 3, 5, 8).

(As I try to redate the exact tree topology of Irisarri et al. [2017], it is not relevant to the present work that interesting doubts about parts of the molecular consensus have been raised from the molecular data, most recently and thoroughly by Mongiardino Koch and Gauthier [2018], who also reviewed that issue.)

The oldest known toxicoferans appear to be represented by four isolated vertebral centra from the Anoual Fm of Morocco, which is early Bathonian in age (Haddoumi et al., 2015). These bones were assigned to “cf. *Parviraptor*” by Haddoumi et al. (2015). Other material – vertebrae and jaw fragments from Europe and North America discussed in Panciroli et al. (2020) – was originally assigned to “cf.” or “aff. *Parviraptor*”, including but not limited to the late Bathonian or earliest Callovian *Eophis*, the Kimmeridgian *Diablophis* and *Portugalophis*, and *Parviraptor* itself from around the Jurassic/Cretaceous (Tithonian/Berriasian) boundary. Traditionally regarded as representing the oldest anguimorphs, these fossils would calibrate Node 130, the split between Pan-Iguania [PN] and Anguimorpha; however, phylogenetic analyses following a redescription of much of the material have found it to constitute the oldest known pan-serpents, thus calibrating Node 129 (Caldwell et al., 2015; Martill et al., 2015; by implication Conrad, 2017; accepted without analysis by Garberoglio et al., 2019, Simões et al., 2020, and Schineider Fachini et al., 2020). As the Bathonian began 168.3 ± 1.3 Ma ago and ended 166.1 ± 1.2 Ma ago, i.e. with uncertainty ranges that overlap in the middle (ICSC), the suggestion of 167 Ma by Caldwell et al. (2015) would then be a reasonable minimum age for this calibration.

Alifanov’s (2019) casual referral of *Parviraptor* to an unusually large version of Mosasauria should not be construed to contradict this: the Cretaceous aquatic squamates, mosasaurs included, are probably all pan-serpents (see below), unless they lie on the common stem of Anguimorpha and Iguania (Simões et al., 2020: supp. fig. 8, with very low support).

As mentioned, all these remains are very fragmentary, and all are disarticulated; according to a reviewer, new, apparently unpublished material shows the “parviraptorids” are not snakes, and indeed Panciroli et al. (2020) were careful not to state in the text whether they agreed with the referral to the snake stem, designating “cf. *Parviraptor* sp.” as “Squamata indet.” in their faunal list (table 1).

The next younger record of a possible toxicoferan is the just as fragmentary Callovian *Changetisaurus*, a supposed anguimorph, though Alifanov (2019) provided reasons to doubt that it is a toxicoferan. It is followed by the several species of *Dorsetisaurus*, another assemblage of skull fragments with osteoderms from the Kimmeridgian through Berriasian of Europe and North America, that was explicitly accepted as an anguimorph by Caldwell et al. (2015) and, on different grounds, Alifanov (2019), but has not, to the best of my knowledge, been included in any phylogenetic analysis. (Older and secondary literature has often claimed that the oldest *Dorsetisaurus* specimens are 148 Ma old, but the Kimmeridgian ended 152.1 ± 0.9 Ma ago: ICSC.)

Most of the rich record of Cretaceous aquatic squamates has traditionally been referred to Anguimorpha, but more likely belongs to Pan-Serpentes (e.g. Garberoglio et al., 2019; Palci et al., 2019; Sobral et al., 2020: fig. S10; Simões et al., 2020: supp. fig. 3, 4, 6, 9; and references therein). It sets in in what seems to be the Hauterivian with *Kaganaias* (Evans et al., 2006; Campbell Mekarski et al., 2019); the Hauterivian ended ∼ 129.4 Ma ago (ICSC, uncertainty not quantified). If neither the “parviraptorids” nor *Changetisaurus* nor *Dorsetisaurus* are accepted as toxicoferans, the minimum age of Node 129 should thus be 130 Ma. To err on the side of caution, that is the age I have used here.

Due to Carroll’s Gap (Marjanović and Laurin, 2013a) I agree with Irisarri et al. (2017) in not assigning a maximum age other than that for Node 125.

#### 2.2.14 Node 131: Iguania [PN] (Pan-Acrodonta [PN] – Pan-Iguanidae [PN])

The origin of Iguania by cladogenesis into Pan-Acrodonta and Pan-Iguanidae was assigned a minimum age of 165 Ma (late Middle Jurassic) and a maximum age of 230 Ma (Carnian, Late Triassic) following Noonan and Chippindale (2006).

*Tikiguania* was described as a Late Triassic acrodontan [PN]. Not only is it an acrodontan, it is a draconine agamid (Hutchinson et al., 2012); most likely, therefore, the very well preserved isolated lower jaw is not a fossil, but belongs to one of the draconine species that live on the site, and fell into the screenwashing sample (Hutchinson et al., 2012).

*Bharatagama*, cited by Noonan and Chippindale (2006), is known (Evans et al., 2002) from at least 85 maxilla and dentary fragments (with supposed genuine absence of the splenial and supposed fusion of the angular to the dentary) that undoubtedly come from the Upper Member of the Kota Fm in Andhra Pradesh (India), for which, on the balance of conflicting biostratigraphic evidence (Prasad and Manhas, 2007; Prasad et al., 2014), a late Middle Jurassic age seems most likely (notwithstanding the fact that the Lower Member conformably overlies the Dharmaram Fm, which extends down into the Triassic as shown by its phytosaurs and aëtosaurs: Goswami et al., 2016). Even so, this age (i.e. 163.5 ± 1.0 Ma or older: ICSC) is old enough by comparison to the pan-iguanian fossil record and the position of Iguania in all molecular phylogenies (including Irisarri et al., 2017) that Jones et al. (2013: 15), whose molecular dating found Toxicofera as a whole to be younger than *Bharatagama*, stated: “It is possible that *Bharatagama* represents an early stem crown-group [sic] squamate with a jaw morphology convergent with modern acrodont [= acrodontan] iguanians, or that it belongs to another clade.” Simões et al. (2017) cited these doubts without further comment. Evans et al. (2002: 306) listed a number of features shared by acrodontans and sphenodontians; three of these do not occur in the Cretaceous priscagamid stem-pan-acrodontans, but all are found in *Bharatagama*. Although no known sphenodontian is a good match (Evans et al., 2002), I very tentatively suggest that *Bharatagama* could represent a morphologically innovative clade of *Diphydontosaurus*-grade sphenodontians. It would not lie outside the large (Reynoso, 2005, and references therein) sphenodontian morphospace: the shape, size, implantation and attachment of the distal teeth recalls *Clevosaurus* (depicted in Evans et al., 2002), while the shape and size of the mesial teeth is reminiscent of *Sphenovipera* (Reynoso, 2005). Indeed, the one phylogenetic analysis that has ever included *Bharatagama* found it as a rhynchocephalian rather than a squamate, although close to the pleurosaurs (despite the more *Diphydontosaurus*-like plesiomorphic gradient of tooth implantation) and, not surprisingly given the limited material, with weak support (Conrad, 2017). In sum, the optimism of Scarpetta (2019) about the usefulness of *Bharatagama* as a calibration point is unwarranted, because the status of *Bharatagama* as a pan-acrodontan is too doubtful.

*Xianglong* from the Yixian Fm of Liaoning (China), which dates to around the Barremian-Aptian boundary (∼ 125.0 Ma: ICSC), was described as a pan-acrodontan, possibly an acrodontan (Li et al., 2007). Unfortunately, this rests on very limited evidence: the one known individual is clearly juvenile, and much of the skeleton remains unknown because is covered by exquisitely preserved soft CT-scanned (Li et al., 2007; Simões et al., 2017; Scarpetta, 2019, and μ reference therein).

Daza et al. (2016) briefly described three isolated hindlimbs from Burmese amber (99 Ma old: Daza et al., 2016, 2020) as agamids, and a largely complete articulated individual as a chamaeleonid. The supposed chamaeleonid later turned out to be an albanerpetid amphibian with a ballistic tongue (Matsumoto and Evans, 2018: 52–53; Daza et al., 2020), and the supposed agamids are so incomplete that they probably provide more ecological than phylogenetic information; indeed, the only supposed pan-acrodontan Daza et al. (2016) included in their phylogenetic analysis was the albanerpetid.

Therefore, again unlike Scarpetta (2019), I do not think any of these four specimens can be used to calibrate divergence dates.

Priscagamidae is a Campanian clade (from the Djadokhta, Baruungoyot and more or less coeval formations; see node 113 above and Borsuk-Białynicka, 1996) of squamates that have usually been considered stem-pan-acrodontans (most recently found as such by Simões et al., 2018, and the three matrices independently derived from theirs: Garberoglio et al., 2019; Sobral et al., 2020; Simões et al., 2020; also by DeMar et al., 2017), but have also been found as stem-pan-iguanians (Conrad, 2015: fig. 6, with much denser sampling of pan-iguanians than in DeMar et al., 2017, or Simões et al., 2018, and their successors).

A consensus now appears to exist that Gobiguania (Conrad and Norell, 2007) is a clade or grade of Campanian and Maastrichtian stem-pan-iguanians (Simões et al., 2015; Conrad, 2015), though DeMar et al. (2017: supp. inf.) could not determine if their two gobiguanian clades were stem-pan-iguanians or stem-pan-iguanids [PN].

“*Ctenomastax*” Gao and Norell, 2000, a junior homonym of the staphylinid beetle *Ctenomastax* Kraatz in von Heyden, 1870, is likewise known from the Djadokhta and Baruungoyot formations (see node 113); probably due to the poor preservation of the specimens (Gao and Norell, 2000), it has variously been found as the sister-group of all other pan-acrodontans (Simões et al., 2015; Reeder et al., 2015; DeMar et al., 2017) or as a gobiguanian stem-pan-iguanian (Conrad, 2015). In the latter case it cannot date the origin of Iguania.

*Isodontosaurus*, from the Djadokhta Fm and more or less coeval sites, is known from fairly large amounts of material representing much of the skeleton, but its phylogenetic position has been hard to determine (Gao and Norell, 2000); Conrad (2015) found it as a stem-pan-acrodontan, Reeder et al. (2015) as a gobiguanian, DeMar et al. (2017) in the “gobiguanian” grade.

DeMar et al. (2017: supp. inf.: 26–28) briefly reviewed the various Cretaceous specimens from North and South America that had been attributed to Pan-Iguanidae [PN], in some cases even Iguanidae [PN] (see node 132), and found all these attributions doubtful at best.

Alifanov (2013) described *Desertiguana* as a phrynosomatid iguanid [PN] based on an almost complete left lower jaw from the Baruungoyot Fm. Curiously, it has been summarily ignored ever since by everyone other than its author (in single-authored publications that do not provide further information and never contain phylogenetic analyses), except for a citation as a pan-iguanian without any comment by Head (2015). Given that Alifanov (2013) also classified three other Djadokhta/Baruungoyot genera otherwise considered gobiguanians as phrynosomatids, I cannot be certain that *Desertiguana* is not a gobiguanian stem-pan-iguanian as well.

Equally Campanian or older (summarized in Langer et al., 2019) is the stem-pan-acrodontan *Gueragama* (Simões et al., 2015, 2017). Known from an isolated but largely complete lower jaw, it appears to suffice for setting up a minimum age for Iguania at the Campanian/Maastrichtian boundary (72.1 ± 0.2 Ma: ICSC), which I round to 72 Ma. I should mention, however, that a reviewer doubts the phylogenetic position of *Gueragama* for unstated reasons, and that Romo de Vivar et al. (2020) found that most or all of the similarities between *Gueragama* and Acrodonta are shared with the Triassic pan-lepidosaur *Cargninia*, likely indicating that these features are evolutionarily correlated with each other and prone to convergence. Meanwhile, Alifanov (2020) called *Gueragama* an isodontosaurid (see above) without stating a reason.

Apesteguía et al. (2016) described *Jeddaherdan* from a Cenomanian jaw fragment. Using a dataset entirely restricted to iguanians, their parsimony analysis recovered it as a pan-acrodontan rather than a pan-iguanid (the only other option) and did not resolve it further until implied weighting was applied, which placed *Jeddaherdan* in a clade with *Gueragama* and the extant agamid *Uromastyx*. Bayesian inference found the same result, although with rather low support (posterior probability of 0.8). As the authors pointed out, this topology implies that the occurrence of tooth replacement in *Gueragama* is a reversal. Given the very limited material, the taxon sample which presupposes that *Jeddaherdan* is an iguanian, the constraints on the applicability of implied weighting and the poorly understood performance of Bayesian inference with missing data distributed by body part (Marjanović and Laurin, 2019, and references therein; King, 2019), as well as the implications for *Gueragam* I prefer not to use *Jeddaherdan* to date the origin of Iguania as long as further material has not been discovered.

If none of the taxa listed above are iguanians, the fossil record of Iguania is entirely restricted to the Cenozoic, possibly beginning in the Thanetian, the last stage of the Paleocene (reviewed in Alifanov, 2020 – a work that is, however, perfectly happy to name paraphyletic taxa that are not intended as clades). I cannot assign a maximum age other than that for Node 125.

#### 2.2.15 Node 132: Iguanidae [PN] (Iguaninae + Corytophanidae – Dactyloidae + Phrynosomatidae)

The origin of Iguanidae was given a minimum age of 125 Ma (Barremian/Aptian boundary, Early Cretaceous) and a maximum age of 180 Ma (Toarcian, Early Jurassic). This was miscopied from Noonan and Chippindale (2006), who did assign a maximum age of 180 Ma, but a minimum age of only 25 Ma (late Oligocene), citing an early Miocene specimen and its description from 1991.

Other than the abovementioned Cretaceous and Paleocene questionable iguanids like *Desertiguana* (see node 131), it is unexpectedly hard to determine from the literature what the oldest possible iguanid could be (though even the questionable ones are all much younger than 125 Ma). Smith (2009) described two assemblages of isolated skull bones from the Paleocene-Eocene boundary (56.0 Ma ago: ICSC) as the new taxa *Suzanniwana*, which he considered a likely stem-corytophanid, and *Anolbanolis*, which he thought close to *Polychrus* and Dactyloidae. He did not perform a phylogenetic analysis. Unfortunately, nobody has ever included *Anolbanolis* in a phylogenetic analysis to the best of my knowledge. DeMar et al. (2017) mentioned it in the text as one of the two oldest definitive iguanids (the other being the younger *Afairiguana*), but it does not occur in their tree figure or their entire supplementary information; *Suzanniwana* occurs nowhere in that publication at all. Conrad (2015), nowhere mentioning *Anolbanolis*, stated that *Suzanniwana* was one of the two “taxa with the most volatile positions within this analysis”, but only published the Adams consensus of that analysis, which shows *Suzanniwana* as part of a polytomy that also encompasses Corytophanidae and a clade containing all other extant iguanids – whether *Suzanniwana* remains inside Iguanidae in all of the 98 most parsimonious trees or is placed as the sister-group of Iguanidae in some could only be determined by repeating the analysis. Scarpetta (2020: supp. inf.) did include *Suzanniwana* in one of the two datasets he analyzed, and found it in the corytophanid total group or at least in a clade with Corytophanidae, *Polychrus* and Dactyloidae, but the sample of extinct species is extremely small in that matrix, and *Anolbanolis* is nowhere mentioned.

The oldest certain iguanid, then, is the oldest one known from articulated remains: the fairly highly nested *Kopidosaurus*, even though it is not clear where it is nested exactly (Scarpetta, 2020). Being slightly older than a 52.59 ± 0.12 Ma old tuff that overlies it (Scarpetta, 2020), and being followed by *Afairiguana* (which forms an exclusive clade with the extant *Polychrus* and Dactyloidae in the analysis of Conrad, 2015), the highly nested corytophanid *Babibasiliscus* and the less highly nested corytophanid *Geiseltaliellus* (Conrad, 2015) within the next five million years, it establishes a rather tight minimum age of 53 Ma for this calibration point, very close to the abovementioned 56 Ma.

If *Desertiguana* is not an iguanid, the absence of iguanids might suggest a late Campanian maximum age for Iguanidae. But as this possibility cannot be excluded at present, even apart from unknown geographic or ecological factors that could have kept iguanids out of the environments that deposited the Campanian and Maastrichtian formations of Asia and North America, I find myself unable to assign a maximum age other than, again, that for Node 125. The argument by Noonan and Chippindale (2006: table 1) was “based on observations of Evans et al. (2002) and the assumption that the origin of this group does not predate the earliest known Iguaninan [sic] in the Jurassic” and is therefore doubly untenable.

Burbrink et al. (2020) found extremely short internal branch lengths for the basal radiation of Iguanidae; similarly, Scarpetta (2020) found the phylogeny of Iguanidae difficult to resolve, which likewise suggests a fast radiation (but might also be a consequence of the sparse taxon sampling in both matrices). Paleoecologically, the recovery phase immediately after the Cretaceous-Paleogene boundary suggests itself as the time of such a radiation. But this remains to be tested.

#### 2.2.16 Node 150: Mammalia [PN] (Pan-Monotremata [PN] – Theriimorpha)

The origin of the crown-group Mammalia by the divergence of Pan-Monotremata represented by *Ornithorhynchus*, on one side, and Theriimorpha, which comprises Theria (to which all extant mammals except the monotremes belong), Spalacotheroidea, Meridiolestida, Dryolestidae, Multituberculata, (Eu)triconodonta and many others, on the other side, was assigned a minimum age of 162.5 Ma (Oxfordian, Late Jurassic) and a maximum age of 191.4 Ma (Early Jurassic) following Benton and Donoghue (2007).

The phylogenetic position of Haramiyida, a clade that reaches beyond these ages into the Late Triassic, has been controversial; Celik and Phillips (2020) have presented a strong argument that it lies well outside Mammalia, which is one of the two positions found in previous analyses.

The oldest uncontroversial mammals are the pan-monotremes *Asfaltomylos* and *Henosferus* and the volaticotherian (eu)triconodont *Argentoconodon*, which all come from a level that was originally thought to be high in the Cañadón Asfalto Fm and to be Callovian or even Oxfordian (late Middle or early Late Jurassic) in age, but has more recently been correlated to its very bottom, the transition with the underlying Lonco Trapial Fm (Cúneo et al., 2013). From this bottom of the Cañadón Asfalto Fm, three successive (from lowest to highest) U-Pb zircon dates were determined by Cúneo et al. (2013): 178.766 ± 0.23 Ma, 177.37 ± 0.12 Ma and 176.15 ± 0.24 Ma. These are maximum ages in that reworked zircon crystals occur in these lacustrine tuff beds, so that the youngest crystals, from which the cited ages were derived, could still be older than the deposition of the tuff beds themselves; however, given the correlation of the recovered ages with stratigraphic height, and the rarity of older zircons in the oldest and the youngest sample (Cúneo et al., 2013), a large discrepancy is unlikely. Therefore, I recommend a minimum age of 179 Ma for this calibration.

The maximum age assigned by Irisarri et al. (2017) may be intended to represent the Sinemurian/Pliensbachian boundary (190.8 ± 1.0 Ma: ICSC). Indeed, the Sinemurian record of mammalomorphs (tritylodontids, tritheledontids, *Sinoconodon*, morganucodontans, *Hadrocodium*) from North America, southern Africa and China is fairly rich and diverse, but has not yielded mammals so far. However, ghost lineages encompassing almost the entire Early Jurassic to the middle of the Middle Jurassic occur for haramiyidans and docodonts, both of which have been found in the Rhaetian and the Bathonian, but not so far in between; and while the Rhaetian and/or possibly Norian *Thomasia* and *Haramiyavia* lie outside the smallest clade of all other haramiyidans, the Rhaetian *Tikitherium* is the sister-group of all Jurassic docodonts except the probably Middle Jurassic *Gondtherium* (Zhou et al., 2019: supp. inf. M), requiring two such ghost lineages within Docodonta. Two more such ghost lineages for Pan-Monotremata and Theriimorpha would not be very surprising.

This may be especially relevant if Haramiyida, rather than the Sinemurian *Hadrocodium*, is the sister-group of Mammalia. Currently, the former is recovered by parsimony, the latter by Bayesian analysis of the same matrix (Huttenlocker et al., 2018: extended data fig. 9; Zhou et al., 2019: supp. inf. M), neither option having strong support by its own criteria; judging from the dashes in their fig. 2 and S1, Celik and Phillips (2020) may have found the same result using an improved version of the same matrix, but they did not publish their most parsimonious trees. For comparisons between the methods as applied to paleontological datasets, see the references cited under node 102 (above). Preferring to err on the side of caution, I place the hard maximum age in the Carnian Pluvial Episode 233 Ma ago (Maron et al., 2018), which is also substantially older than all possible haramiyidans, indeed older than all currently recognized mammalomorphs (Kligman et al., 2020, and references therein).

#### 2.2.17 Node 151: Theria (Metatheria – Eutheria)

The origin of Theria by the split into the total groups Metatheria (crown group: Marsupialia) and Eutheria (crown group: Placentalia) was given a minimum age of 124.6 Ma (Barremian/Aptian boundary, Early Cretaceous) and a maximum age of 138.4 Ma (Valanginian, Early Cretaceous) following Benton and Donoghue (2007).

The oldest securely dated therian is currently the stem-eutherian *Ambolestes* at 126 Ma (Bi et al., 2018).

*Juramaia* (Luo et al., 2011) has often been cited as a much older stem-eutherian. However, both its age and its phylogenetic position are in doubt; if either of these doubts is corroborated, *Juramaia* becomes irrelevant to dating this node. Originally, the only known specimen was thought to come from the Lanqi Fm, specifically a site variably called Daxigou or Daxishan (Yuan et al., 2013: supp. inf.: 4), which has meanwhile been dated to between 160.889 ± 0.069 Ma and 160.254 ± 0.045 Ma (Jia and Gao, 2019). Meng (2014: 526, 529–530), however, doubted this, called the specimen “floating”, and pointed out its great similarity to *Eomaia* in particular (found as its sister-group in the very different matrices of Bi et al., 2018, and Zhou et al., 2019: supp. inf. M; Mao et al., 2019: fig. S9, did find *Juramaia* outside the clade of all other included eutherians, but did not sample *Ambolestes* despite building on the matrix of Bi et al., 2018) and to Barremian–Albian eutherians in general, as well as the long ghost lineages a mid-Oxfordian age for *Juramaia* would create within Eutheria, for Metatheria and for several of the closest relatives of Theria. Bi et al. (2018, 2019) referred to Meng (2014) for this issue but did not try to resolve it. As long as it is not resolved, I much prefer to consider the single *Juramaia* specimen to have been discovered in the Yixian Fm (like *Ambolestes*, *Eomaia* and *Acristatherium*), as suggested by Bi et al. (2019).

Celik and Phillips (2020) called *Juramaia* “purportedly Jurassic” without comment and found middling support for a sister-group relationship to Theria as a whole, noting that this agreed with earlier doubts (e.g. by Sweetman et al., 2017). However, like Mao et al. (2019), they did not sample *Ambolestes*, and the sensitivity of this result to whether parsimony or a model-based method is used was not published.

Sweetman et al. (2017) described two teeth from the very beginning of the Cretaceous (∼ 145 Ma old) as two genera of Late-Cretaceous-grade eutherians, *Durlstotherium* and *Durlstodon*. In view of this limited material, I remain skeptical (see also Bi et al., 2018) and recommend 126 Ma as the minimum age for this calibration.

While the oldest uncontested metatherians are only some 110 Ma old (Bi et al., 2018), Mao et al. (2019: fig. S9) and Celik and Phillips (2020) have returned *Sinodelphys* (of the same age as *Eomaia* and *Acristatherium*, slightly younger than *Ambolestes*) to its status as the oldest known metatherian.

If this holds and if *Juramaia* has the same age instead of being Jurassic or is not a therian, and if further *Durlstotherium* and *Durlstodon* can be disregarded, virtually no ghost lineage is required at the base of Metatheria.

Accepting that *Juramaia* is not from the Lanqi Fm or not a therian, I propose 160 Ma as the soft maximum age of this calibration, on the grounds that therians or their closest relatives – other than, perhaps, *Juramaia* – are absent in the Lanqi Fm and the laterally equivalent Tiaojishan Fm, likewise absent in the Kimmeridgian and Tithonian of Portugal and the US (where the Morrison Fm, intensely sampled since the 1860s, extends across several states), and further absent in the end-Tithonian and Berriasian of England – other than, perhaps, *Durlstotherium* and *Durlstodon* – despite the diversity of ecologically comparable mammals found there. Given the strong evidence of a Laurasian origin of Theria (e.g. Huttenlocker et al., 2018; Bi et al., 2018), the earliest possible time and place for the origin of Theria that could stay out of the fossil record is therefore Asia after the deposition of the Tiaojishan and Lanqi formations ended in the Oxfordian.

#### 2.2.18 Node 152: Placentalia (Atlantogenata – Boreo(eu)theria); Node 153: Boreo(eu)theria (Laurasiatheria – Euarchontoglires/Supraprimates)

The origin of Placentalia, the crown group of Eutheria, was given a minimum age of 95.3 Ma (Cenomanian, Late Cretaceous) and a maximum age of 113 Ma (Aptian/Albian boundary, Early Cretaceous) following Benton and Donoghue (2007). Its immediate descendant nodes were not constrained.

The minimum age rests on the assumption, commonly but not universally held in 2007, that the zhelestids are “ungulates”, i.e. belong to Placentalia, or perhaps even that the zalambdalestids are related to Glires and therefore belong to Placentalia. For a long time now, as already pointed out by Parham et al. (2011), every reinvestigation of the anatomy of these Cretaceous animals, and every phylogenetic analysis that sampled Cretaceous eutherians densely (i.e. not including Zhou et al., 2019: supp. inf. M), has found them on the eutherian stem, often not even particularly close to Placentalia (e.g. Novacek et al., 1997; Asher et al., 2005, 2019; Wible et al., 2009; Goswami et al., 2011; Halliday et al., 2015; Manz et al., 2015; Bi et al., 2018: fig. 2, SI-1; Wang et al., 2019: ext. data fig. 5; and references in Parham et al., 2011 “2012”; see also Fostowicz-Frelik and Kielan-Jaworowska, 2002).

A few terminal Cretaceous (late Maastrichtian) eutherians have been attributed to Placentalia in the past. This is at best dubious for all of them. *Protungulatum* (Wible et al., 2009; Halliday et al., 2015, 2019: fig. 1 contrary to the text; Manz et al., 2015: fig. 2a; Wang et al., 2019: ext. data fig. 5; Mao et al., 2019: fig. S9) and *Gypsonictops* (Halliday et al., 2015, 2019; Manz et al., 2015: fig. 2; Bi et al., 2018; Wang et al., 2019: ext. data fig. 5; Mao et al., 2019: fig. S9) are now placed close to but consistently outside Placentalia. *Deccanolestes* – at least if the teeth and the tarsal bones belong together – is placed far away (Goswami et al., 2011 [see there also for *Sahnitherium*]; Manz et al., 2015: fig. 2, SI-1; Penkrot and Zack, 2016; Halliday et al., 2019). The single worn tooth named *Kharmerungulatum*, which had been assigned to Placentalia mostly through comparison to *Protungulatum* in the first place (Prasad et al., 2007), has more recently been found outside Placentalia as well (“Although none of the strict consensus trees supported the placement of *Kharmerungulatum* within the placental crown group, the limited dental material for this taxon proved insufficient for resolving its phylogenetic relationships, and so it was removed a posteriori from the MPTs to produce the reduced strict consensus trees.” – Goswami et al., 2011: 16334), specifically as an adapisoriculid like *Deccanolestes* when full molecular constraints were applied by Manz et al. (2015: fig. 2b). The stylinodontid taeniodont *Schowalteria* (Fox, 2016, and references therein) belongs to a clade that survived into the Eocene; the conference abstract by Funston et al. (2020) reported that a very large phylogenetic analysis has found the group outside Placentalia.

The same reasons make it difficult to decide which of the earliest Paleocene eutherians should be accepted as securely enough identified placentals. But in any case, Williamson et al. (2019: 220) reported that the herbivorous periptychid *Ectoconus*, estimated to have reached about 100 kg, was “present within 0.4 Ma of the K-Pg boundary”; phylogenetic analyses have found it to be not only a placental, but a laurasiatherian – Halliday et al. (2015; regardless of constraints) found it and the other periptychids on the pholidotan stem; Halliday et al. (2019), using combined data and maximum likelihood, found a comparable result with much less resolution; Püschel et al. (2019), using a somewhat smaller matrix with, however, a focus on periptychids and new data on them, recovered them as stem-artiodactylomorphs. I therefore suggest 66 Ma, the Cretaceous/Paleogene boundary (66.021 ± 0.081 Ma: Clyde et al., 2016), as the minimum age for Node 153, the basal node of Boreoeutheria (a name apparently coined by accident by Murphy et al., 2001) or simply Boreotheria (explicitly coined by Waddell et al., 2001). For Node 152 I cannot recommend a separate minimum age.

Unambiguous placentals continue to be absent worldwide in the rich Maastrichtian record (see above as well as Halliday et al., 2016, and Davies et al., 2017), and even ambiguous ones except *Gypsonictops* continue to be absent in the even richer Campanian record (although there are three isolated Turonian teeth indistinguishable from both species of *Gypsonictops*: Cohen and Cifelli, 2015; Cohen, 2017), despite the presence of stem-eutherians (all northern continents, Madagascar and India), stem-metatherians (Asia and North America), and ecologically comparable spalacotheroids (Asia and North America), meridiolestidans (South America) and gondwanatheres (South America, Madagascar, India, and some point between the late Turonian and latest Campanian of Africa – O’Connor et al., 2019). Thus, only Antarctica, Australia and New Zealand are left as paleocontinents where Campanian or Maastrichtian placentals could have escaped the fossil record, and they are all unlikely for biogeographical reasons (e.g. Huttenlocker et al., 2018). Therefore, I suggest the Campanian/Maastrichtian boundary, rounded to 72 Ma, as the hard maximum age for Node 152. (I cannot make a separate recommendation for Node 153.) This is more generous than the result of Halliday et al. (2016), 95% of whose reconstructions of the age of Placentalia were 69.53 Ma old or younger. The discrepancy to the published molecular ages (references in Halliday et al., 2016) is most likely due to the effects of body size (Berv and Field, 2017; Phillips and Fruciano, 2018), or perhaps other factors like generation length, on rates of molecular evolution.

At this point, readers may be wondering why I have mentioned neither the extremely large phylogenetic analysis by O’Leary et al. (2013) nor the objections by Springer et al. (2019), who wrote in their abstract that “morphological cladistics has a poor track record of reconstructing higher-level relationships among the orders of placental mammals”. It would be more accurate to say that phylogenetic analysis of morphological data has *no* track record of reconstructing the phylogeny of Placentalia, good *or* bad. To avoid long-branch attraction and long-branch repulsion, any such analysis of morphological data will have to sample the enormous and poorly understood diversity of Paleo- and Eocene eutherians very densely, which will have to entail sampling enough of the characters that unite and distinguish them without falling into the trap of accumulating redundant or otherwise correlated characters that inevitably distort the tree (Marjanović and Laurin, 2019; Sookias, 2019; Celik and Phillips, 2020; and references in all three). This is so much work, and so hard to get funded, that at the most generous count only three attempts at such a matrix have ever been made; I should also point out that matrices of such sizes were not computationally tractable until a few years ago, at least not in less than a few months of calculation time. The first attempt is the “phenomic” matrix by O’Leary et al. (2013); as Springer et al. (2019) pointed out repeatedly, it contains no less than 4,541 characters – but several hundred of these are parsimony-uninformative (O’Leary et al., 2013), and many others are redundant, which means they represent a smaller number of independent characters of which many are weighted twice or more often. At 86 terminal taxa, almost all of which are extant, the taxon sample is hopelessly inadequate for eutherian phylogeny. It is no surprise that parts of the topology are highly implausible (e.g. the undisputed stem-whale *Rodhocetus* landing on the common ungulate [PN] stem, as pointed out by Springer et al., 2019) and that even such undisputed clades as Afrosoricida, Lipotyphla and Artiodactyla are no longer recovered when the hundreds of soft-tissue characters, which cannot be scored for the extinct terminal taxa, are removed (Springer et al., 2019), which casts doubt on the ability of that matrix to place extinct taxa accurately. The second attempt began in the doctoral thesis of Zack (2009) and was further modified and merged with other datasets in Halliday’s doctoral thesis that culminated in the publication of Halliday et al. (2015). The taxon sample contains an appreciable number of Cretaceous and Paleocene eutherians; the character sample is of course more modest and contains, as usual for mammals, a large proportion of tooth characters, some of which might be redundant (e.g. Kangas et al., 2004; Harjunmaa et al., 2014). The further improved version (Halliday et al., 2019) suffers from the drawback that all characters were reduced to two states to make the matrix tractable by maximum-likelihood software; this throws away a lot of information (probably for no gain: Sansom et al., 2018; King, 2019). The third is that of the PalM group; funded by an enormous grant, it involves a lot of people each revising a group of Paleo- or Eocene eutherians as their doctoral thesis and contributing the gained knowledge (e.g. Napoli et al., 2017) to a growing matrix (ultimately based on that of Wible et al., 2009) that will then be evaluated for character redundancy and other issues. The only phylogenetic publications that have yet resulted are conference abstracts, of which I have cited Püschel et al. (2019) and Funston et al. (2020) above.

Springer et al. (2019) went on to claim that “Sansom and Wills (2013) showed that fossils are more likely to move stemward than crownward when they are only known for biomineralized characters”. Indeed Sansom and Wills (2013) made that claim. They had taken 78 neontological matrices of extant animals with biomineralized tissues, deleted the data for soft-tissue characters from random taxa and found that those taxa changed their phylogenetic position significantly more often than random, and further underwent “stemward slippage” as opposed to “crownward slippage” significantly more often than random. Deleting data from hard-tissue characters instead had no such effect. Sansom and Wills (2013) concluded that some mysterious factor causes hard-tissue characters to contain a systematically misleading signal much more often than soft-tissue characters do, and that therefore the phylogenetic positions of all taxa known only from hard tissues – in other words most animal fossils – are highly suspect of falsely appearing more rootward than they really are. Therefore, fossils assigned to various stem groups could really belong to the crown groups, and the minimum ages of divergence-date calibrations could be systematically too young (Sansom and Wills, 2013), just as Springer et al. (2019) believed. A much simpler explanation is available: hard-tissue characters are unreliable *specifically among extant species* because the hard-tissue anatomy of extant species is usually very poorly known. For example (Marjanović and Witzmann, 2015), the vertebrae of some of western and central Europe’s most common newt species are simply unknown to science, even after 200 years or more of research, because neontologists have focused on soft-tissue anatomy, behavior and more recently the genome while treating the skeleton as an afterthought. And the vertebrae of salamandrids are at least known to contain a phylogenetic signal – whether the appendicular skeleton also does is anybody’s guess at this point! As our knowledge of the skeletons of extant taxa would improve, so would, I predict, the ability of hard-tissue characters to accurately resolve the phylogenetic positions of extant taxa.

#### 2.2.19 Node 154: Carnivora [PN] (Pan-Feliformia [PN] – Pan-Caniformia [PN])

The origin of Carnivora by the divergence of the sister-groups Pan-Feliformia (represented in this matrix by *Felis*) and Pan-Caniformia (represented by *Canis*) was assigned a minimum age of 42.8 Ma (Lutetian, Eocene) and a maximum age of 63.8 Ma (Danian, Paleocene). Irisarri et al. (2017) justified this by citing the identification of the middle Eocene *Tapocyon* as a pan-caniform by Wesley and Flynn (2003); this should be regarded as rendered obsolete by Spaulding and Flynn (2012) and Solé et al. (2016), who found *Tapocyon* as a stem-carnivoriform in phylogenetic analyses of two successively larger versions of a much larger dataset. The analysis by Tomiya and Tseng (2016) found *Tapocyon* as a pan-feliform, but used a much smaller sample of stem-carnivoriforms and of characters in a misguided (e.g. Kearney and Clark, 2003; Wiens, 2003a, b, 2005a, b; Prevosti and Chemisquy, 2009; Marjanović and Laurin, 2019; King, 2019; Mongiardino Koch et al., 2020) attempt to avoid missing data by throwing out known data.

With “*Miacis*” *sylvestris* being recovered even more rootward on the carnivoriform stem than *Tapocyon* by Spaulding and Flynn (2012) and Solé et al. (2016), the oldest securely dated and identified carnivoran specimens belong to the amphicyonid stem-pan-caniform *Daphoenus* and the stem-canid *Hesperocyon* and are about 38 Ma old (Tomiya, 2011, and references therein). *Lycophocyon* could have the same age or be somewhat older (Tomiya, 2011), but unfortunately its phylogenetic position remains uncertain: it was published too late to be included by Spaulding and Flynn (2012), it was not added by Solé et al. (2016), and the much smaller phylogenetic analysis by Tomiya (2011) only resolved its position (as a stem-pan-caniform closer to Caniformia than *Daphoenus*) after all post-Paleogene taxa were excluded. Given the uncertainties in both age and phylogenetic position, I provisionally ignore *Lycophocyon* and suggest 38 Ma as the minimum age of this calibration.

As a hard maximum age I suggest the Paleocene/Eocene boundary 56.0 Ma ago (ICSC), around which there is a very rich record of a range of carnivorous mammals of various sizes and ecologies, including stem-carnivoriforms and many others but not including carnivorans.

#### 2.2.20 Node 155: Euarchontoglires/Supraprimates (Gliriformes – Primatomorpha)

The last common ancestor of mice and men, the first crown-group member of a clade called Euarchontoglires (a name apparently coined by accident by Murphy et al., 2001) or, perhaps less clunkily, Supraprimates (explicitly coined by Waddell et al., 2001), was placed between 61.5 Ma ago (Selandian, Paleocene) and 100.5 Ma ago (Early/Late Cretaceous boundary) following Benton and Donoghue (2007).

The oldest purported total-group primatomorph – not necessarily a pan-primate [PN] (Ni et al., 2016) – is *Purgatorius coracis*, found in an outcrop of the Ravenscrag Formation that is at most 0.4 Ma younger than the 66.0-Ma-old Cretaceous/Paleogene boundary (Fox and Scott, 2011; Scott et al., 2016). However, Halliday et al. (2015, 2019) found *Purgatorius* outside of Placentalia despite the presence of stem-pan-primates in their analyses. When Manz et al. (2015) applied molecular constraints (fig. 2), they did find *Purgatorius* as a pan-primate, though in a strangely nested position when the monophyly of Laurasiatheria was enforced (fig. 2b). Without constraints, the included primatomorphs formed a grade outside most other placentals (and the included laurasiatherians formed a grade outside all other placentals: fig. SI3-1). Note that Halliday et al. (2015, 2019) scored *Purgatorius* for the tarsal bones that Chester et al. (2015) referred to this taxon (somewhat younger than *P. coracis*); *Purgatorius* is otherwise known exclusively from teeth and lower jaws (Chester et al., 2015; Scott et al., 2016), and Chester et al. (2015) referred the tarsals simply because their size fits and because they show arboreal adaptations which agree with the assumed pan-primate status of *Purgatorius*. Scott et al. (2016: 343) preferred to call these bones “several isolated, possible plesiadapiform tarsals”, Plesiadapiformes being a clade or grade of stem-pan-primates or stem-primatomorphs to which *Purgatorius* is generally thought to belong.

Excluding the purgatoriids, the diverse oldest known total-group primatomorphs are, in terms of North American Land Mammal Ages, slightly younger than the Puercan/Torrejonian boundary (Silcox et al., 2017), which dates to about 64.8 Ma ago (Wang et al., 2016).

On the presumably gliriform side, the oldest known members are anagalidans from the Lower Member of the Wanghudun Fm: the anagalids *Anaptogale*, *Wanogale* and *Chianshania*, the pseudictopid *Cartictops* and the astigalid *Astigale* (Missiaen et al., 2012; Wang et al., 2016; López-Torres and Fostowicz-Frelik, 2018). Their ages are poorly constrained between 66 Ma and about 62.5 Ma, though probably closer to the older end of that range (Wang et al., 2016); López-Torres and Fostowicz-Frelik (2018: fig. 4) illustrated *Anaptogale* as considerably older than *Wanogale* and *Chianshania*, but did not explain why. However, Asher et al. (2019: fig. 4, S5B, supplementary file S4-optimalTrees.nex) found Anagalida in a “primatomorph grade” when using equally weighted parsimony or implied weights with K = 24, as afrotherians with K = 2, and on the eutherian stem by Bayesian inference; at least in the latter two cases, anagalidans cannot calibrate this node.

Thus, I propose 65 Ma as the minimum age of this calibration. As the maximum age, if 66 Ma is deemed too close to the minimum (although there are presently no proposed crown- or even total-group supraprimates from the Cretaceous, despite the abundance of ecologically Glires-like and early-primatomorph-like multituberculates, gondwanatheres and the presence – in India – of ecologically pan-primate-like adapisoriculids) or to the age of the oldest *Purgatorius*, I can only offer the maximum of Node 152 (Placentalia, see above).

#### 2.2.21 Node 157: Marsupialia (Didelphimorphia – Paucituberculata + Australidelphia)

The origin of the metatherian crown group Marsupialia was given a minimum age of 61.5 Ma (Selandian, Paleocene) and a maximum age of 71.2 Ma (Maastrichtian, Late Cretaceous) following Benton and Donoghue (2007).

Eldridge et al. (2019) reviewed this question, and found that the oldest definitive marsupials are only 54.6 Ma old as far as understood today, dating from shortly after the beginning of the Eocene (56.0 Ma ago: ICSC). Their phylogenetic and geographic position (total-group australidelphians from Australia) suggests a longer history for Marsupialia, but of the many metatherians known from the Paleocene of South America and from the Late Cretaceous through Miocene of the northern continents, none can currently be shown to belong to the crown group (Eldridge et al., 2019). I therefore propose 55 Ma as a probably overly strict minimum age for this calibration.

Carneiro (2017; not cited by Eldridge et al., 2019, whose paper was accepted for publication on 15 January 2018) found the Maastrichtian tooth taxon *Glasbius* from North America as a didelphimorphian marsupial in a phylogenetic analysis (greatly expanded from that of Carneiro and Oliveira, 2017, with the same result, likewise not cited by Eldridge et al., 2019). That analysis, however, implied an extraordinary number of transoceanic dispersals around the Paleocene and – as the Gondwanan metatherians are all Cenozoic, but most Laurasian ones are Mesozoic – a surprisingly high rate of survival of metatherians across the Cretaceous/Paleogene boundary. I must suspect that correlation, if not downright redundancy, among mammalian tooth characters has been underestimated once again (e.g. Kangas et al., 2004; Harjunmaa et al., 2014; Celik and Phillips, 2020). Indeed, Cohen et al. (2020b) found *Glasbius* on the metatherian stem; however, although they discussed this result, they did not cite Carneiro (2017) or Carneiro and Oliveira (2017). Their analysis also failed to find the two included australidelphian taxa as sister-groups despite the morphological and molecular consensus (see Eldridge et al., 2019), but the bootstrap support for this was low.

Marsupials, other metatherians and indeed other therians are wholly absent from the Late Cretaceous mammaliform record of South America, which consists instead of gondwanatherian haramiyidans, a few multituberculates and a very wide variety of meridiolestidan stem-theriiforms. The ages of the latest Cretaceous terrestrial sites of South America have been difficult to pinpoint, but there is evidence that they cover the entire Campanian and Maastrichtian (Rougier et al., 2008; Lawver et al., 2011; and references therein). The early Paleocene (Danian) sites of South America do contain stem-metatherians (and eutherians; references in Eldridge et al., 2019). If *Glasbius* is not a marsupial, it can be stated with great confidence that Marsupialia originated in South America (Eldridge et al., 2019, and references therein); if *Glasbius* is a marsupial, North America becomes the obvious candidate, and at least two clades of marsupials most likely survived the Cretaceous and immigrated into South America separately. In that case, it is noteworthy that *Glasbius* is the only possible marsupial out of the remarkable diversity of Maastrichtian, Campanian and in some cases yet earlier metatherians known from North America and to a lesser degree central Asia. Rather than the beginning of the Maastrichtian, I propose the beginning of deposition of the Lance and Hell Creek formations, where *Glasbius* has been found, as the hard maximum age for this calibration, which I estimate as 68 Ma – though the single tooth from the Williams Fork Fm that Cohen et al. (2020b) referred to *Glasbius* may be up to 2 Ma older.

#### 2.2.22 Node 160: Batrachia (Urodela – Salientia)

The origin of Batrachia by the divergence of the sister-groups Urodela (the salamander total group now that Caudata [PN] is the crown group) and Salientia (the frog total group) was assigned a minimum age of 249 Ma and no maximum age. This was, as usual, done on the basis of *Triadobatrachus*, one of the two oldest known salientians (the other is *Czatkobatrachus*, which is probably early Olenëkian in age: Evans and Borsuk-Białynicka, 2009); all known definitive urodeles are considerably younger (Schoch et al., 2020). Irisarri et al. (2017) only cited the classic redescription of *Triadobatrachus* from 1989 for this age; more recent stratigraphic work has been reviewed by Ascarrunz et al. (2016: 206–207) and places *Triadobatrachus* either in the late Induan or the very beginning of the Olenëkian. Unfortunately, the precise age of the Induan-Olenëkian boundary remains unclear; the ICSC, indirectly citing a source from 2007, places it at 251.2 Ma without explicit error margins, while Maron et al. (2018) placed it at “∼ 249.7 Ma” based on cyclostratigraphic counting away from the Permian-Triassic boundary, which is exceptionally precisely dated radiometrically. I conclude that 249 Ma is a perfectly adequate minimum age for this calibration point.

For a maximum age, I reiterate the suggestion of Marjanović and Laurin (2013b) to use the beginning of Carroll’s Gap (see Marjanović and Laurin, 2013a), i.e. the Early Permian record, which has yielded many tetrapods ecologically comparable to batrachians, but no batrachians, gymnophionomorphs or albanerpetids so far (e.g. Schoch and Milner, 2014; Glienke, 2015). The abovementioned particularly rich site of Richards Spur, where small terrestrial and possibly amphibious temnospondyls particularly similar to batrachians are very common, has yielded three radiometric ages, of which the oldest is 289.2 ± 0.68 Ma old (Woodhead et al., 2010; MacDougall et al., 2017), so that 290 Ma may be a defensible soft maximum value. (The value of 275 Ma suggested by Marjanović and Laurin, 2007 and 2013b, is outdated.)

#### 2.2.23 Node 169: crown group of Cryptobranchoidea (Hynobiidae – Pancryptobrancha)

The divergence between the salamander clades Pancryptobrancha (the smallest total group containing the crown group Cryptobranchidae: Vasilyan et al., 2013) and Hynobiidae was assigned a minimum age of 145.5 Ma and no maximum age.

The minimum age, intended to correspond to the Jurassic/Cretaceous boundary (∼ 145.0 Ma ago: ICSC), constitutes a snapshot in the convoluted history of dating the Jurassic and Cretaceous Konservat-Lagerstätten of northeastern China. (Another such snapshot, likewise outdated, is the Valanginian age of 139.4 Ma suggested for this node by Marjanović and Laurin, 2007.) None of these sites are now considered Kimmeridgian through Valanginian in age. The oldest ones that have yielded purported caudates [PN] (references in Skutschas, 2015, beginning with Gao and Shubin, 2003, the reference cited for this calibration by Irisarri et al., 2017) belong to the Daohugou Beds, which correlate with the Haifanggou Fm and are Callovian (late Middle Jurassic) or earliest Oxfordian (Late Jurassic) in age (Gao and Shubin, 2012; Jiang et al., 2015; Liang et al., 2019; Rong et al., accepted; and references therein), not Bathonian as often claimed in older literature. These lithostratigraphic units immediately underlie the abovementioned (see node 151) Lanqi and Tiaojishan formations, which have – including in the abovementioned Daxishan or Daxigou site – likewise yielded purported caudates (Gao and Shubin, 2012; Jia and Gao, 2016, 2019).

Two Bathonian sites with supposed crown-group salamanders do exist. One (Kirtlington, UK; Forest Marble Fm) has yielded at least one undescribed vertebra called “Kirtlington salamander B”. The other (Berezovsk, Russia; Itat Fm) has yielded *Kiyatriton krasnolutskii* Skutschas, 2015; while the association of the isolated bones from different body parts with each other is rather daring, the holotype of this species (like the holotype of the much younger type species, another isolated atlantal centrum) does preserve a clear synapomorphy with Caudata and three similarities to Cryptobranchoidea (Skutschas, 2014, 2015). Both sites have also yielded isolated femora that show one potential synapomorphy with Hynobiidae (Skutschas, 2014, 2015). Potentially, then, *K. krasnolutskii* could be the oldest known crown-cryptobranchoid and necessitate a minimum age of about 169 Ma (ICSC) for this node. Unfortunately, no bone referred to *Kiyatriton* has yet been included in a phylogenetic analysis, and that is not likely to happen soon: the two existing morphological datasets for analysis of salamander phylogeny (latest published versions: Wiens et al., 2005; Rong et al., accepted) are very light on atlas characters, which moreover are mostly not accessible in the Chinese Mesozoic specimens (complete, articulated, flattened skeletons with soft-tissue outlines and melanosomes) and not well understood in extant salamanders – like the rest of the skeleton in general and the postcranial skeleton in particular, which neontologists have by and large ignored in favor of molecular, behavioral and soft-tissue characters (see Marjanović and Witzmann, 2015, for some drastic examples).

The latest published phylogenetic analysis of Mesozoic salamanders is that by Rong et al. (accepted). Like the morphological subset of Wiens et al. (2005), it produces – unless a molecular constraint is applied – a clear example of what Wiens et al. (2005: title) called “[o]ntogeny discombobulates phylogeny”: a clade composed of the extant neotenic non-cryptobranchoid salamander clades, i.e. (Amphiumidae (Sirenidae, Proteidae)), as the sister-group of the metamorphic ones. Indeed, its character sample is full of characters that translate straightforwardly to presence vs. absence of a complete metamorphosis (or of a strictly aquatic lifestyle). (That is in addition to simpler, even more objective problems in the character list of the lineage of matrices from Gao and Shubin [2012] through Jia and Gao [2016, 2019] to Rong et al. [accepted]; for example, in all four of these matrices, characters 77 and 78 are duplicates of each other – the haploid number of chromosomes and the diploid number of chromosomes.) Instead, molecular data (e.g. Wiens et al., 2005; Irisarri et al., 2017; Vijayakumar et al., 2019: supplementary file Amphibia_New_India_SHL_Dryad.tre; Hime et al., 2020; and references therein) have consistently shown that Sirenidae lies outside the smallest clade formed by all other extant non-cryptobranchoid salamanders (Salamandroidea), as had long been presumed based on other considerations like the retention of external fertilization in sirenids (Reinhard et al., 2013). Likewise, Amphiumidae and Plethodontidae are consistently sister-groups in phylogenetic analyses of molecular data, rather than Amphiumidae being close to Proteidae or Sirenidae, or Plethodontidae being close to Salamandridae (e.g. Rong, 2018; Rong et al., accepted) or *Ambystoma* (e.g. Jia and Gao, 2019). This may be particularly relevant because all of the Chinese Mesozoic salamanders are either only known from larval or neotenic individuals (e.g. *Chunerpeton*: Rong et al., accepted), or are metamorphic but aquatic (*Pangerpeton*: Wang and Evans, 2006), or combine features expected of different ontogenetic stages (perhaps indicating a less condensed metamorphosis than in extant metamorphic salamanders: *Linglongtriton* [Jia and Gao, 2019]; also *Chunerpeton* [Rong et al., accepted] and, though found outside Cryptobranchoidea, *Beiyanerpeton*: Gao and Shubin, 2012), or are metamorphic and apparently terrestrial but have not been sufficiently described to be included in a phylogenetic analysis (*Laccotriton*). All known possible pancryptobranchans except the terminal Paleocene stem-pancryptobranchan *Aviturus* (Vasilyan and Böhme, 2012; Vasilyan et al., 2013) have been neotenic or undergone only partial metamorphosis (the extant *Andrias* loses the gills, the gill slits and the coronoid bone, but does not rebuild the palate or grow eyelids); this may attract stem-cryptobranchoids or even some of the more crownward stem-urodeles toward them, even if some (Rong, 2018) or most (Jia and Gao, 2019) or a variable number (Rong et al., accepted) end up in the hynobiid total group rather than in Pancryptobrancha. Unfortunately, no published phylogenetic analysis has ever included extinct Cenozoic pancryptobranchans together with any Mesozoic salamanders; the overlap between the taxon samples of Vasilyan et al. (2013) and Pearson (2016) or Rong et al. (accepted), as well as all references in all three, is restricted to extant species.

I should point out that plesiomorphies unexpected in caudates have been found in some of the Chinese Mesozoic taxa. For example, as pointed out by Marjanović and Laurin (2019: appendix S1: 76) and confirmed by Rong et al. (accepted), free palatines occur in *Chunerpeton* (Wang et al., 2015; illustrated in Gao and Shubin, 2003, though not indicated or mentioned in the text), *Beiyanerpeton* (Gao and Shubin, 2012) and *Qinglongtriton* (Jia and Gao, 2016). This appears to be borne out by the phylogenetic analyses of Rong et al. (accepted).

It does not help that the known fossil record of possible hynobiids outside of the mentioned Lagerstätten only begins in the late Miocene and consists entirely of isolated bones (reviewed by Jia and Gao, 2016: 44–45). One possible exception is the metamorphic *Iridotriton*, known from a partial but well preserved skeleton from the early Tithonian (Galli et al., 2018; Maidment and Muxworthy, 2019) Brushy Basin Member of the Morrison Fm (Rainbow Park Microsite, Utah), originally argued to be a non-cryptobranchoid caudate (Evans et al., 2005), more recently found in an incompletely resolved position outside the cryptobranchoid crown-group (Pearson, 2016: fig. 4.11; Rong et al., accepted), and equipped with a confusing combination of characters (Evans et al., 2005).

Mesozoic pancryptobranchans seem to be represented by a large number of isolated bones from the early Cenomanian through early Campanian of Kazakhstan, Uzbekistan and Tajikistan (Skutschas, 2013) usually grouped as *Eoscapherpeton* and *Horezmia* (but considered two species of *Eoscapherpeton* by Skutschas, 2013). Unfortunately, they have never been included in a phylogenetic analysis outside of Pearson’s (2016) doctoral thesis, but the arguments by Skutschas (2013) for referring at least some of the nonoverlapping material to Pancryptobrancha are not easily dismissed. In a Bayesian analysis of a matrix of morphological data containing extant lissamphibians, the Permian temnospondyls *Doleserpeton* and *Gerobatrachus*, the stem-salientian *Triadobatrachus*, *Eoscapherpeton* and a number of Cretaceous and Cenozoic scapherpetids but no other caudates, Pearson (2016: fig. 4.2) recovered *Eoscapherpeton* as a stem-pancryptobranchan, though with a posterior probability of only 52%; adding further Mesozoic salamanders led to the breakdown of this resolution (Pearson, 2016: fig. 4.12).

The oldest wholly undoubted pancryptobranchan is “*Cryptobranchus*” *saskatchewanensis*, which has been included in the phylogenetic analysis of Vasilyan et al. (2013). It comes from an exposure of the same Ravenscrag Fm that is mentioned under Node 155, but widely separated in space and age from the one mentioned there: in terms of North American Land Mammal Ages, the site with the oldest “*C.*” *saskatchewanensis* specimens – including the holotype – is Tiffanian-4 in age, thus between 59 and 60 Ma (Krause, 1978; Naylor, 1981; Wang et al., 2016: fig. 2). The material consists of isolated dentary fragments (like the holotype), maxilla fragments and an exoccipital referred by size alone; they all seem to be within the morphological range of known (Cenozoic) pancryptobranchans, but not more convincing than the similarly fragmentary *Eoscapherpeton*.

I therefore use the beginning of the Cenomanian (100.5 Ma ago, given without uncertainty in the ICSC), rounded to 101 Ma ago, as the minimum age of this calibration for present purposes. Given the great uncertainty, I generally recommend against using this divergence as a calibration.

(My previous suggestion – Marjanović, 2019 – to use this age as a soft minimum was incoherent, as a reviewer pointed out. A soft minimum would imply that a tail of the probability distribution of the age of this node would extend to younger ages than 101 Ma, so that an age of 100 Ma would be treated as much more probable than an age of, say, 61 Ma. The opposite is the case: both 101 and 60 are much more probable than 100, which is younger than one potential minimum age but far older than the other. If *Eoscapherpeton* is a crown-group cryptobranchoid, so that 101 Ma is “the correct” minimum age, 100 is impossible; if it is not a crown-group cryptobranchoid, so that 60 is “correct”, 100 is so much older as to be much less probable than, say, 65.)

It is interesting in this respect that calibrating this node with an age around 139.4 Ma (Marjanović and Laurin, 2007) leads to far too high ages for cladogeneses within Hynobiidae and within Cryptobranchidae, even within *Andrias japonicus* judging by paleogeographic criteria (Matsui et al., 2008).

Like Irisarri et al. (2017), I cannot assign a maximum age other than that of Node 160. The oldest known stem-salamanders, except for the Middle or Late Triassic *Triassurus* (Schoch et al., 2020), are Bathonian (Skutschas, 2015, and references therein); the fossil record of total-group salamanders thus exemplifies Carroll’s Gap (Marjanović and Laurin, 2013a).

#### 2.2.24 Node 170: Lalagobatrachia/Bombinanura (total group of Bombinatoroidea/Costata – total group of Pipanura); Node 171: Pipanura (total group of Pipoidea/Xenanura – total group of Acosmanura)

The last common ancestor of Bombinatoroidea or Costata, represented by *Bombina* and *Discoglossus*, and Pipanura, to which all other sampled frogs belong, was assigned a minimum age of 161.2 Ma (Oxfordian, Late Jurassic) and no maximum age. Pipanura itself was assigned a minimum age of 145.5 Ma (end-Jurassic) and no maximum age.

Following the finding that *Eodiscoglossus*, to which a Bathonian (Middle Jurassic) species has been referred that forms the basis for the original minimum age, is probably not a costatan (Báez, 2013; Báez and Gómez, 2016, 2019), the oldest purported lalagobatrachian/bombinanuran is the poorly known *Enneabatrachus* from a site dated to 152.51 ± 0.47 Ma (Trujillo et al., 2015), which has never been included in a phylogenetic analysis. Given, however, the presence of the pipanuran (rhinophrynid or stem-xenanuran: Henrici, 1998; Gómez, 2016; Aranciaga Rolando et al., 2019) *Rhadinosteus* at the same site as *Iridotriton* (the Rainbow Park Microsite, see node 169) and as further specimens of *Enneabatrachus*, a minimum age of 153 Ma for Pipanura (and Bombinanura by implication), coinciding with the maximum age of the Kimmeridgian/Tithonian boundary (152.1 ± 0.9 Ma: ICSC) and constituting a minimal revision of the age proposed by Marjanović and Laurin (2013b), appears safe.

*Enneabatrachus*, if not *Rhadinosteus*, is at present the oldest securely identified anuran (crown-group salientian). Remarkably, no salientians at all have so far been reported from the Yanliao Biota (Haifanggou, Lanqi, Tiaojishan and maybe other formations of Callovian to Oxfordian age in northeastern China), despite its wealth of salamanders (see node 169). The stem-salientian record is sparse (Marjanović and Laurin, 2013b; Stocker et al., 2019); the suggestion of a maximum age for Bombinanura of 170 to 185 Ma by Marjanović and Laurin (2013b) is based on the fairly good stratigraphic fit of stem-salientian phylogeny (Marjanović and Laurin, 2007, 2013a: fig. 5, 2013b; Stocker et al., 2019; and references therein), but given its poor geographic coverage, I prefer to follow Irisarri et al. (2017) in not assigning a maximum age other than that of node 160 for present purposes.

Thus, node 170 cannot currently be calibrated on its own: its minimum age is that of node 171, its maximum age is that of node 160.

#### 2.2.25 Node 178: Pipidae (Pipinomorpha – Xenopodinomorpha)

The origin of Pipidae (the crown group of Pipimorpha) by the divergence of Pipinomorpha (crown group: Pipinae) and Xenopodinomorpha (crown group: Xenopodinae = *Xenopus* sensu lato) was given a minimum age of 86 Ma (Coniacian/Santonian boundary, Late Cretaceous) and no maximum age.

This cladogenesis is particularly difficult to date from the fossil record because molecular data support Hymenochirini as a member of Xenopodinomorpha, though less strongly than most other parts of the tree (Cannatella, 2015: fig. 1, with a bootstrap support of 71% while other branches have 74%, 93% or 100%, and with a Bayesian posterior probability of 99% while three others have 100%; Irisarri et al., 2017, with a jackknife support of 98% instead of the usual 100%; Vijayakumar et al., 2019: supplementary file Amphibia_New_India_SHL_Dryad.tre, with a Shimodaira/Hasegawa-like approximate likelihood ratio of 91% instead of the usual 100%; Hime et al., 2020: supp. fig. 4, with a bootstrap support of 100% but a local posterior measure of branch support of only 50.77% instead of the usual 80%–100%), while morphological data have so far only supported Hymenochirini as a member of Pipinomorpha (with a Bayesian posterior probability of 100% in Cannatella, 2015). The only phylogenetic analysis of combined data from pipimorphs yet undertaken (Cannatella, 2015: analysis E1) found almost equal support for both possibilities (bootstrap support of 46% vs. 44%; Bayesian posterior probabilities below 50%), and the winning-sites test could not distinguish between them (p = 1.0: Cannatella, 2015: table 3), although tip-dating with three node calibrations strongly supported the hymenochirins as pipinomorphs at the cost of losing a terminal taxon (*Pachycentrata*, see below; Cannatella, 2015: analysis E6).

Using considerably updated and expanded versions of the morphological dataset Cannatella (2015) had used, Gómez (2016), de Souza Carvalho et al. (2019) and Aranciaga Rolando et al. (2019) all found the Cenomanian *Oumtkoutia* (not included by Cannatella, 2015) to be the oldest known pipid; the Cenomanian ended 93.9 Ma ago (ICSC, no error margin given). However, while the first of these three phylogenetic analyses found it as a stem-xenopodinomorph, the other two – whose matrices are almost identical to each other, and derived from that of the first with rather few changes – found it as a stem-pipinomorph, and the third cautioned that it may well be a stem-pipimorph because, although Rage and Dutheil (2008) described the material in great detail, it consists entirely of isolated braincases, vertebrae and pelves, and there is some character conflict as *Oumtkoutia* combines a pipinomorph autapomorphy with stem-pipimorph plesiomorphies. The next younger pipid remains *Pachycentrata* of end-Coniacian or Santonian age, found as a stem-hymenochirinomorph by Gómez (2016) but as a stem-pipinomorph by de Souza Carvalho et al. (2019) and Aranciaga Rolando et al. (2019); while the Coniacian ended 86.3 ± 0.5 Ma ago, the Santonian ended only 83.6 ± 0.2 Ma ago (ICSC).

Given the presence of *Pipa* in South America but its extant sister-group Hymenochirini in Africa, and further the facts that all known pipomorphs are strictly aquatic and that lissamphibians in general tend to tolerate saltwater poorly, it is tempting to assume that this distribution is due to vicariance and the cladogenesis that separated *Pipa* and the hymenochirins should be dated to the loss of contact between Outer Gondwana (including South America) and Afro-Arabia around the Cenomanian – in other words, a geological event should be used to calibrate this divergence date. If *Pachycentrata* is a stem-hymenochirinomorph, as found by Gómez (2016), this scenario fits the phylogeny beautifully, and neither any overseas dispersal nor any long ghost lineages need to be assumed, as Gómez (2016) pointed out. Contrariwise, if *Pachycentrata* is a stem-pipinomorph, as found by de Souza Carvalho et al. (2019) and Aranciaga Rolando et al. (2019), the fossil record offers no reason to date the origin of Pipinae to the Mesozoic, and the most parsimonious hypothesis becomes that *Pipa* dispersed from Africa to South America together with the platyrrhine monkeys and the caviomorph rodents, perhaps on the same natural raft; de Souza Carvalho et al. (2019: 228) have discussed the possibility of a Paleogene island chain or even landbridge on the Walvis Ridge and the Rio Grande Rise at some length.

On the phylogenies by de Souza Carvalho et al. (2019) and Aranciaga Rolando et al. (2019), the xenopodinomorph fossil record begins only in the late Oligocene (briefly reviewed in Blackburn et al., 2019; see also Gardner and Rage, 2016: 184) rather than the Cenomanian (Gómez, 2016).

As mentioned, the only combined dataset yet brought to bear on this question (Cannatella, 2015: dataset E), which is also the only dataset containing extinct taxa that supports the hymenochirins as pipinomorphs, is based on a superseded morphological dataset that lacked *Oumtkoutia* and *Pachycentrata*, not to mention any taxa described since 2007. Given this and the discussion in the preceding paragraphs, it remains unclear whether *Oumtkoutia* is a pipid, and so I can only suggest 84 Ma as a safe minimum age for Pipidae.

Any maximum age will have to accommodate the undescribed possible pipid from the Aptian or Barremian of Cameroon (Gardner and Rage, 2016: 177, 179). However, the only maximum age I feel able to propose is much older: the end of deposition of the lake sediments of the Newark Supergroup (Tanner and Lucas, 2015) sometime around the Hettangian/Sinemurian boundary (199.3 ± 0.3 Ma ago: ICSC). All known pipimorphs, extant or extinct, have been fully aquatic (reviewed in Cannatella, 2015). The upper formations of the Newark Supergroup, which represent the rift lakes that preceded the opening of the Central Atlantic Ocean between Africa and North America, have yielded whole species flocks of semionotid actinopterygians among other parts of a lake fauna and flora (Olsen, 1988, 2010), and they cover so much space and time that if any aquatic salientians existed in northwestern Pangea during that time, we should expect to have found them – yet, salientians are consistently absent from these sediments (Olsen, 1988). The absence of salamanders (Olsen, 1988) may be explained by geography in that that group may have originated in Asia or at least northeastern Pangea (where indeed the Middle or Late Triassic *Triassurus* was found: Schoch et al., 2020). All other Barremian or earlier xenoanurans, however, have so far been found on the Iberian microcontinent or in North America, and the stratigraphic fit of their phylogeny (Gómez, 2016; Aranciaga Rolando et al., 2019) is good enough that if pipids older than *Oumtkoutia* existed, northwestern Pangea is where we should look for them. I therefore propose 199 Ma as the hard maximum age for this calibration.

It may be significant that anurans have not so far been found in the lacustrine Bathonian sediments (∼ 167 Ma old) of the Anoual Fm in Morocco (Haddoumi et al., 2015).

#### 2.2.26 Node 187: crown group of Chondrichthyes (Holocephali – Elasmobranchii)

The origin of the chondrichthyan crown group was given a minimum age of 410 Ma (Lochkovian/Pragian boundary, Devonian) and a maximum age of 495 Ma (Paibian, Furongian, Cambrian). Note that the maximum age was not operational because the root node was given a younger maximum age of 462.5 Ma.

By current understanding (Frey et al., 2019), the oldest known crown-chondrichthyan is the stem-elasmobranch *Phoebodus fastigatus* from the middle Givetian. The Givetian, part of the Middle Devonian, began 387.7 ± 0.8 Ma ago and ended 382.7 ± 1.6 Ma ago (ICSC), so I propose 385 Ma as the minimum age of the chondrichthyan crown-group.

Although I cannot assign a maximum age separate from that of the root node (node 100) to this calibration, no less than ninety million years before the minimum age, I note that this is still twenty million years after the 495 Ma assigned, futilely, by Irisarri et al. (2017).

#### 2.2.27 Node 188: crown group of Elasmobranchii (Selachimorpha – Batomorpha)

The origin of the elasmobranch crown group by split into Selachimorpha (sharks) and Batomorpha (rays and skates) was given a minimum age of 190 Ma (Sinemurian/Pliensbachian boundary, Early Jurassic) and no maximum age. (Note that the name Neoselachii is consistently treated in the paleontological literature as if defined by one or more apomorphies, not by tree topology; it probably applies to a clade somewhat larger, and possibly much older, than its crown group.)

Any attempt to date this cladogenesis suffers from the fact that the elasmobranch fossil record consists mostly of ‘the tooth, the whole tooth and nothing but the tooth’ (as has often been said about the Mesozoic mammalian fossil record); scales and the occasional fin spine do occur, but more substantial remains are very rare. The shape of tooth crowns is naturally prone to homoplasy, the number of phylogenetically informative characters it offers is easily overestimated due to correlations between them (e.g. Kangas et al., 2004; Harjunmaa et al., 2014; Celik and Phillips, 2020; see node 157), and histological studies, which are needed to determine the states of certain characters (e.g. Andreev and Cuny, 2012; Cuny et al., 2017), have not been carried out on all potentially interesting tooth taxa.

Consequently, there is not as much interest in phylogeny among specialists of early elasmobranchs than among specialists of early mammals or early dinosaurs. This goes so far as to affect the use of terminology: Andreev and Cuny (2012) mentioned “stem selachimorphs” in the title of their work, implying that they understood Selachimorpha as a clade name, but quietly revealed it to be the name of a paraphyletic assemblage on p. 263 by stating that bundled enameloid is “diagnostic for Neoselachii exclusive of batomorphs, i.e., Selachimorpha”, and their consistent referral of Synechodontiformes (see below) to “Selachimorpha” is not necessarily a referral to the crown group – even though they called bato- and selachomorphs sister-groups in the next sentence.

A safe minimum age of 201 Ma, used here, is provided by the oldest unambiguous crown-group selachimorph, the total-group galeomorph *Agaleus*, dating from the Hettangian, apparently close to its beginning (Stumpf and Kriwet, 2019, especially fig. 5, and references therein), which was the beginning of the Jurassic and happened 201.3 ± 0.2 Ma ago (ICSC); I round this down (stratigraphically up) to avoid breaching the mass extinction event at the Triassic/Jurassic boundary. The oldest batoid batomorph is only sightly younger, see node 192 below. However, this may err very far on the side of caution. Indeed, for purposes beyond the present work, I must recommend against using the minimum age of this divergence to calibrate a timetree for at least as long as the histology of Paleozoic “shark” teeth has not been studied in much more detail in a phylogenetic context. As if by typographic error, the oldest widely accepted crown-group elasmobranch is not 190 but about 290 Ma old: the oldest fossils referred to the neoselachian *Synechodus* are four teeth of Sakmarian age (referred to *S. antiquus*, whose type tooth comes from the following Artinskian age: Ivanov, 2005; Stumpf and Kriwet, 2019), and the Sakmarian ended 290.1 ± 0.26 Ma ago (ICSC). Teeth referred to other species of *Synechodus* range into the Paleocene; *S. antiquus* is the only Permian species (Andreev and Cuny, 2012). The histology of *S. antiquus* remains unknown as of Koot et al. (2014); nonetheless, Cuny et al. (2017: 61) regarded *S. antiquus* as “[t]he first proven selachimorph”. Rounding up, this would suggest suggest 291 Ma as the minimum age of this calibration.

(My previous suggestion – Marjanović, 2019 – to use that age as a soft minimum was incoherent, as a reviewer pointed out. A soft minimum would imply that a tail of the probability distribution of the age of this node would extend to younger ages than 291 Ma, so that an age of 290 Ma would be treated as much more probable than an age of 201 Ma. The opposite is the case: both 291 and 202 are much more probable than 290, which is younger than one potential minimum age but far older than the other. If *Synechodus antiquus* is a crown-group elasmobranch, so that 291 Ma is “the correct” minimum age, 290 is impossible; if it is not a crown-group elasmobranch, so that 201 is “correct”, 290 is so much older as to be much less probable than, say, 205 or 210.)

Potential crown-group elasmobranchs older than 291 Ma are known: Andreev and Cuny (2012) and Cuny et al. (2017: 69) suggested that the tooth taxa *Cooleyella* and *Ginteria* could be stem-batomorphs. The oldest known *Cooleyella* specimen dates from around the end of the Tournaisian (Richards et al., 2018), which occurred 346.7 ± 0.4 Ma ago (ICSC); *Ginteria* appeared in the following Viséan stage. Cuny et al. (2017: 21, 69) further pointed out that *Mcmurdodus*, a tooth taxon that first appeared around the Early/Middle Devonian (Emsian/Eifelian) boundary (Burrow et al., 2008), has occasionally been placed within Selachimorpha, even within Hexanchiformes in the selachimorph crown-group (Burrow et al., 2008, and references therein); they very tentatively suggested a stem-selachimorph position. Boisvert et al. (2019) wondered instead if it is a stem-chondrichthyan.

The absence of any however tentative suggestions of crown-elasmobranchs before *Mcmurdodus* in the rather rich total-group chondrichthyan microfossil record despite the traditional optimism of paleodontologists may, somewhat ironically, serve as a hard maximum age for this calibration; the ICSC places the Emsian/Eifelian boundary at 393.3 ± 1.2 Ma ago, so I suggest 395 Ma.

#### 2.2.28 Node 192: Batoidea (skates – rays)

The origin of the batomorph crown group, Batoidea, by split into skates (Rajiformes; represented by *Raja* and *Leucoraja*) and rays (taxonomically unnamed; represented by *Neotrygon*) was assigned a minimum age of 176 Ma (Toarcian, Early Jurassic) and no maximum age.

The oldest known batoid is a single rajiform tooth named *Antiquaobatis* from the late Pliensbachian, specifically the *apyrenum* subzone of the *spinatum* ammonite zone (Stumpf and Kriwet, 2019), which is close to the end of the Pliensbachian (Fraguas et al., 2018); that end occurred 182.7 ± 0.7 Ma ago (ICSC), so I propose 184 Ma as the minimum age for this calibration. (The name should of course have been “Antiquobatis”, but must not be amended: ICZN, 1999: Article 32.5.1.)

As a hard maximum age, the Triassic/Jurassic boundary (201.3 ± 0.2 Ma ago: ICSC; rounded to 201 Ma) suggests itself for ecological reasons: plesiomorphically, crown-group rays are fairly large marine durophages, a guild formed by the placodont amniotes in the well sampled Middle and Late Triassic.

#### 2.2.29 Node 195: Neopterygii [PN] (Holosteomorpha – Pan-Teleostei [PN])

The origin of Neopterygii by cladogenesis into the total groups of Holostei (bowfins – *Amia* – and gars, represented by *Lepisosteus*) and Teleostei [PN] (represented by the clupeocephalans *Takifugu* and *Danio*) was given a minimum age of 345 Ma and a maximum age of 392 Ma.

At present, there are only two candidates for Paleozoic neopterygians. One is *Acentrophorus*, “a ‘semionotid’-like taxon that desperately requires restudy and formal analysis” (Friedman, 2015: 222; cited as current by Xu, 2019; also Sun et al., 2016) of Wujiapingian age (between 254.14 ± 0.07 Ma and 259.1 ± 0.5 Ma: ICSC). The “semionotids” are stem-members of Ginglymodi, i.e. closer to *Lepisosteus* than to *Amia* (Giles et al., 2017: ext. data fig. 6–8; López-Arbarello and Sferco, 2018; Xu, 2019), but a generic “‘semionotid’-like taxon” could easily lie outside Neopterygii. In their in-depth study of neopterygian phylogeny, López-Arbarello and Sferco (2018) did not include *Acentrophorus* or even mention it in the text.

Sun et al. (2016) cited *Archaeolepidotus*, supposedly closely related to *Watsonulus* (see below), together with undescribed specimens as a Changxingian neopterygian (which was originally thought to be Early Triassic, but probably is not according to references in Ronchi et al., 2018). The Changxingian is the stage between the Wujiapingian and the Permian/Triassic boundary (251.902 ± 0.024 Ma ago: ICSC). *Archaeolepidotus* does not appear to be well understood; Friedman (2015), Giles et al. (2017), López-Arbarello and Sferco (2018) and Xu (2019) did not mention it, let alone include it in a phylogenetic analysis, and Google Scholar only finds 17 occurrences in the entire literature.

The oldest certain member of Neopterygii is *Watsonulus*, a stem-halecomorph or stem-holosteomorph (Friedman, 2015; Giles et al., 2017: ext. data fig. 6–8; López-Arbarello and Sferco, 2018; Xu, 2019) which comes from the Middle Sakamena Group of Madagascar (López-Arbarello and Sferco, 2018) just like *Triadobatrachus* (see node 160) and should therefore be around 249 Ma old. I therefore propose 249 Ma as the minimum age of Neopterygii.

Assuming from the almost phylogeny-free quantification of the Permo-Triassic fossil record of osteichthyans by Romano et al. (2014b) that at least the Asselian record of pan-actinopterygians [PN] is reasonably good, I suggest a soft maximum age for Neopterygii immediately before it, i.e. at the Carboniferous/Permian boundary (298.9 ± 0.15 Ma: ICSC), rounded to 299 Ma, which conveniently places it 50 Ma before the minimum age.

### 2.3 Analysis methods

Johan Renaudie (Museum für Naturkunde, Berlin) kindly performed the divergence dating using the tree (topology and uncalibrated branch lengths), the model of evolution (CAT-GTR+Γ) and clock model (lognormal autocorrelated relaxed) inferred by Irisarri et al. (2017) and the data (“nuclear test data set”: the variable sites of the 14,352 most complete amino acid positions of their “NoDP” dataset), but the calibrations presented above (all at once, not different subsets).

The intent was to also use the software Irisarri et al. (2017) had used (PhyloBayes, though the latest version, 4.1c: Lartillot, 2015). However, PhyloBayes is unable to treat some bounds as hard and others as soft in the same analysis; it can only treat all as soft, as Irisarri et al. (2017) had done, or all as hard. Consequently, we ran our analysis with all bounds treated as hard in order to account for the hard minima (discussed above: Materials and methods: Hard and soft minima and maxima).

The launch code for our PhyloBayes analysis is: ./pb -d ali14352.phy -T final_tree.tre -cal dm4.txt -r outgroups -bd -cat -gtr -ln -dc dm4hardDC.1 & ./pb -d ali14352.phy -T final_tree.tre -cal dm4.txt -r outgroups -bd -cat -gtr -ln -dc dm4hardDC.2

Irisarri et al. (2017) ran 100 gene-jackknifed analyses for each of their two sets of calibrations. Lacking the necessary computational resources, we only ran two analyses of the full dataset, without jackknifing. The results (Table 2, Fig. 1) are therefore less reliable, given the data, than those of Irisarri et al. (2017); but they fully suffice as a proof of concept to show that improved calibrations lead to many different inferred node ages.

Above I describe phylogenetic uncertainty leading to two different minimum ages for Tetrapoda (node 105), 335 Ma and “roughly” 350 Ma. Using the younger age results in a younger bound of 359 Ma on the 95% credibility interval of this node (mean age: 363 Ma, older bound: 365 Ma, i.e. the maximum age of the calibration: Table 2); therefore, I do not consider it necessary to set the minimum age of this node to 350 Ma and run a second analysis.

## 3 Results and discussion

### 3.1 Bibliometry

Irisarri et al. (2017: supp. table 8) cited 15 works as sources for their calibrations, six of them compilations made by paleontologists to help molecular biologists calibrate timetrees.

Not counting Irisarri et al. (2017) and the ICSC (which has been updated at least once a year since 2008), I cite 235 references to discuss minimum ages (mostly for the age or phylogenetic position of a potentially calibrating specimen), 26 to discuss maximum ages (mostly to argue if observed absence of a clade is reliable), and 15 for both purposes. Of the total of 276, one each dates to 1964, 1978, 1981, 1988 and 1991, 2 each to 1994, 1995 and 1996, 1 each to 1997 and 1998, 3 to 1999, 1 to 2000, 2 to 2001, 4 to 2002, 1 to 2003, 0 to 2004, 7 to 2005, 4 to 2006, 6 each to 2007 and 2008, 4 to 2009, 5 to 2010, 8 to 2011, 9 to 2012, 15 to 2013, 12 to 2014, 23 to 2015, 24 to 2016, 23 to 2017, 28 to 2018, 50 to 2019, 27 to 2020, and one was published as an accepted manuscript in 2020 and may come out this or next year in final form. (Whenever applicable, these are the years of actual publication, i.e. public availability of the layouted and proofread work, not the year of intended publication which can be a year earlier, and not the year of print which is very often one or even two years later.) Only three of these are among the 14 used by Irisarri et al. (2017), and none of them are among the six compilations they cited.

Irisarri et al. submitted their manuscript on 16 September 2016. Assuming that half of the publications cited here that were published in 2016 came out too late to be used by Irisarri et al. (2017), the total proportion of the works cited here that would have been useful to them for calibrating their timetree but were not available amounts to 140 of 276, or 50.7%. Similarly, 249 of the works cited here, or 90.2%, were published since mid-2005. I conclude from this extreme “pull of the recent” that knowledge in this area has an extremely short half-life; calibration dates, therefore, cannot be taken from published compilations (including the present work) or other secondary sources, but must be checked every time anew against the current primary literature. This is time-consuming even in the digital age, much more so than I expected, and requires reading more works for context than actually end up cited (for some nodes three times as many); but there is no shortcut.

### 3.2 Changes in the calibration dates

Of the 30 minimum ages assigned by Irisarri et al. (2017), I find only one to be accurate by the current state of knowledge, that of Batrachia (node 160) anchored by good old *Triadobatrachus* (see Ascarrunz et al., 2016, for the latest and most thorough redescription and stratigraphy, and Daza et al., 2020, for the latest and largest phylogenetic analysis).

The minimum age of Pleurodira (node 124), which has long been known to be 100 Ma older than Irisarri et al. (2017) set it, turns out to be copied from the calibration of a much smaller clade in Noonan and Chippindale (2006), a secondary source whose minimum age for Pleurodira was actually better by a factor of four. The minimum age of Iguanidae (node 132) turned out to be miscopied, most likely with a typographic error, from Noonan and Chippindale (2006), who had it as 25 Ma instead of the 125 Ma of Irisarri et al. (2017) – though 25 Ma is not tenable either, but too young by at least 28 Ma.

In four more cases (Osteichthyes: node 102; Reptilia: node 107; Placentalia: node 152; Lalagobatrachia/Bombinanura: node 170) I find myself unable to assign any minimum age specific to that node. In two of these cases (Reptilia and Placentalia) the specimen previously thought to constrain that node actually constrains a less inclusive clade (Archelosauria, node 108; Boreo(eu)theria, node 153) that was sampled but not constrained by Irisarri et al. (2017); I have used these minimum ages to constrain the latter two nodes.

As might be expected, 15 of the minimum ages are too young, by margins ranging from 1.4 Ma to 100 Ma or, ignoring Pleurodira, 43.25 Ma (Table 1: last two columns). Unsurprisingly, this also holds for the two nodes that Irisarri et al. (2017) did not calibrate but I did: both of them were constrained by calibrated nodes whose minimum ages were too young for these two nodes. In eight cases, including Boreo(eu)theria (node 153), the reason is the expected one, the more or less recent discovery of previously unknown fossils (mostly before 2016); the magnitude of the resulting changes ranges from 1.4 Ma to 11 Ma. In four more cases, including the one used by Irisarri et al. (2017) to date Osteichthyes (node 102) but by me to date the subsequent split of Dipnomorpha and Tetrapodomorpha (node 104), the dating of the oldest known specimens has improved by 4 to 16.5 Ma. The specimen used to constrain Tetrapoda (node 105) is probably not a tetrapod, but the oldest known certain tetrapods are now nonetheless dated as roughly 5 Ma older than the minimum assigned by Irisarri et al. (2017); depending on the phylogenetic hypothesis, isolated bones or (!) footprints roughly 20 Ma older that were published in 2015 could represent the oldest tetrapods instead. The remaining six cases, including Reptilia (node 107) and Archelosauria (node 108) by implication, are caused by phylogenetic reassignments of previously known specimens (mostly before 2016) and have effects ranging from 4 Ma to 43.25 Ma.

The minimum ages of the remaining 13 nodes (including, accidentally, Iguanidae) are too old; the margins vary from 1 Ma to 96 Ma. This includes the case of Toxicofera (node 129), whose minimum age of 148 Ma assigned by Irisarri et al. (2017) was not operational as that node was in fact constrained by the minimum age of its constituent clade Iguania (node 131), 165 Ma; both of these ages are too old – I find minimum ages of 130 Ma for Toxicofera and 72 Ma for Iguania. Interestingly, none of the changes to minimum ages are due to more precise dating. There is one case of the opposite: I have changed the minimum age of Pipidae (node 178) from 86 to 84 Ma because the oldest known safely identified pipid, *Pachycentrata*, may be somewhat older than the Coniacian/Santonian boundary (86.3 ± 0.5 Ma ago: ICSC), but also somewhat younger, so the Santonian/Campanian boundary (83.6 ± 0.2 Ma ago: ICSC) is a safer approximation. All others are due to more or less recent findings that the oldest supposed members of the clades in question cannot, or at least cannot be confidently, assigned to these clades.

I agree with the reasoning for one of the maximum ages used by Irisarri et al. (2017), that for Archosauria (node 109), though its numeric value had to be increased by 1 Ma due to improved dating of the Permian/Triassic boundary since the source Irisarri et al. (2017) used was published in 2005.

I find myself unable to assign a separate maximum age to seven of the 18 remaining nodes that Irisarri et al. (2017) used maximum ages for; these nodes are only constrained by the maximum ages of more inclusive clades in my reanalysis. This includes the case of Chondrichthyes (node 187), whose maximum age of 495 Ma assigned by Irisarri et al. (2017) was not operational as that node was in fact constrained by the maximum age of the root node, 462.5 Ma; I can likewise constrain it only by the maximum age of the root, 475 Ma. In one of these cases the new implied maximum age is younger (by 28.5 Ma) than the previously explicit maximum; in the remainder it is older by 27 Ma to 110 Ma.

Of the remaining 11 maximum ages, six were too young by 12.5 Ma to 125 Ma. In one case (the root: Gnathostomata, node 100), the old maximum is younger than the new minimum, and in two more cases (Mammalia and Theria), phylogenetic (or, in the case of Theria, possibly stratigraphic) uncertainty is the reason; the remaining three merely show greater caution on my part in interpreting absence of evidence as evidence of absence.

The remaining five I consider too old by 3.2 Ma to 93 Ma; these show greater confidence on my part in interpreting absence of evidence as evidence of absence in well-sampled parts of the fossil record. The same holds, naturally, for the six nodes that lacked maximum ages in Irisarri et al. (2017) but that I propose maximum ages for; one of these new ages, however (for Lepidosauria, node 125), is older than the previously implied maximum age provided by the next more inclusive clade, and that by 33 Ma. The other five are 60.1 Ma to no less than 261.5 Ma younger than their previously implied equivalents.

### 3.3 Changes in the divergence dates

Reanalyzing the data of Irisarri et al. (2017) with their methods, but using the calibration ages proposed and discussed above and treating them all as hard bounds in PhyloBayes instead of treating all as soft (see Materials and methods: Hard and soft minima and maxima, Analysis methods), generally leads to implausibly old ages and large credibility intervals for the unconstrained nodes (Fig. 1, Table 2): e.g., the last common ancestor of chickens and turkeys (node 115) is placed around the Cretaceous/Paleogene boundary, with a 95% credibility interval that spans half of each period, and the credibility interval of the bird crown-group (Aves, node 112) spans most of the Jurassic, with a younger bound less than 10 Ma younger than the age of the distant stem-avialan *Archaeopteryx* (just over 150 Ma), while the oldest known crown-birds are less than half as old, about 71 Ma (see Materials and methods: Calibrations: Node 113).

There are exceptions, however. Most notably, the squamate radiation (nodes 126–129) is constrained only between the origin of Lepidosauria (see above under node 125: 244–290 Ma ago) and the origin of Toxicofera (see above under node 129: minimum age 130 Ma), yet it is bunched up close to the latter date, unlike in Irisarri et al. (2017) where it was more spread out and generally older even though both calibrations were younger. For example, the unconstrained origin of Squamata (node 126) was found to have a mean age of 199 Ma by Irisarri et al. (2017), but 153 Ma here (Table 2). The crucial difference may be that Lepidosauria did not have a maximum age, but this does not explain the very short internodes from Squamata to Iguania in my results. I should point out that the oldest likely squamate remains are close to 170 Ma old (reviewed in Panciroli et al., 2020). In part, these implausible ages may be due to effects of body size (Berv and Field, 2017) or loosely related factors like generation length: most sampled squamates are small, while the two sampled palaeognath birds (node 116, with an evidently spurious mean age of 163 Ma) are much larger than all sampled neognaths. This may be supported by the body size increase in snakes: their oldest sampled node (Macrostomata or Afrophidia: node 136) is placed around the Early/Late Cretaceous boundary, followed by the origin of Endoglyptodonta (node 138) in the Late Cretaceous, while any Late Cretaceous caenophidians (a clade containing Endoglyptodonta) remain unknown, all potential Cretaceous total-group macrostomates are beset with phylogenetic uncertainty, and considerably younger dates were found by Burbrink et al. (2020) despite the use of a mid-Cretaceous potential macrostomate as a minimum-age-only calibration. Similarly, the fact that the entire credibility interval for Supraprimates/Euarchontoglires (node 155) was younger than its calibrated minimum age when all bounds were treated as soft in Marjanović (2019) may be due to the fact that one of the two sampled supraprimates is *Homo*, the second-largest sampled mammal and the one with the second-longest generation span.

Whelan and Halanych (2016) found that the CAT-GTR model (at least as implemented in PhyloBayes) is prone to inferring inaccurate branch lengths, especially in large datasets; this may turn out to be another cause of the results described above. The omission of the constant characters from the dataset, intended to speed up calculations (Irisarri et al., 2017), may have exacerbated this problem by leading to inaccurate model parameters (Whelan and Halanych, 2016).

It is, however, noteworthy that all terminal branches inferred here are longer, in terms of time, than in Irisarri et al. (2017).

Naturally, the changes to the calibration dates have changed the inferred ages of many calibrated nodes and the sizes of their credibility intervals. For instance, Irisarri et al. (2017) inferred a mean age of 207 Ma for Batoidea, with a 90-Ma-long 95% credibility interval that stretched from 172 Ma ago to 262 Ma ago (node 192; Table 2); that node was calibrated with a soft minimum age set to 176 Ma, but not only was no maximum age set, no other node between there and the root node (Gnathostomata, node 100) had a maximum age either, so that effectively the maximum age for Batoidea was that of the root node, 462.5 Ma. Following the discovery of new fossils, I have increased the hard minimum age to 184 Ma; however, out of ecological considerations, I have also introduced a hard maximum age of 201 Ma, younger than the previously inferred mean age. Naturally, the new inferred mean age is also younger: 193 Ma, with a 95% credibility interval that spans the time between the calibration dates (Table 2).

Somewhat similarly, I have increased the minimum age of Mammalia (node 150) from 162.5 to 179 Ma following improved dating of the oldest certain mammals, increased its maximum age from 191.4 Ma to 233 Ma to account for phylogenetic uncertainty and the limits of the Norian (middle Late Triassic) fossil record, and treated both bounds as hard. While Irisarri et al. (2017) found a mean age of 165 Ma with a credibility interval from 161 Ma to 172 Ma, straddling the minimum age but not reaching the maximum, I find an age range that reaches the new maximum but stays far away from the new minimum (mean: 229 Ma, 95% credibility interval from 217 Ma to 233 Ma). While the next less inclusive calibrated node (151: Theria) has an increased maximum but a barely changed minimum age, both bounds of the next more inclusive calibrated node (106: Amniota) have increased by about 30 Ma, apparently pulling the inferred age of Mammalia with them.

### 3.4 Pitfalls in interpreting the descriptive paleontological literature

It is widely thought that paleontologists are particularly eager to publish their specimens as the oldest known record of some taxon. Indeed it happens that five different species of different ages are published as the oldest record of the same taxon within ten years. In such cases, finding a specimen that can establish a minimum age for that taxon can be as simple as finding the latest publication that makes such a claim; and that can be as simple as a Google Scholar search restricted to the last few years. But there are harder cases; I will present two.

Above (Materials and methods: Calibrations: Node 132 – Iguanidae) I argue for using the age of *Kopidosaurus*, about 53 million years, as the minimum age of Iguanidae. *Kopidosaurus* was named and described from a largely complete skull by Scarpetta (2020) in a publication where the words “oldest” and “older” do not occur at all, and “first” and “ancient” only occur in other contexts – even though Scarpetta (2019) had just published on calibration dates for molecular divergence date analyses. The reason may be that he did not think *Kopidosaurus* was the oldest iguanid; one of the two matrices he used for phylogenetic analyses contained the 56-Ma-old *Suzanniwana*, and his analyses found it as an iguanid (Scarpetta, 2020: supp. inf.). Moreover, he was most likely aware that the publication that named and described *Suzanniwana* (Smith, 2009) also named and described *Anolbanolis* from the same site and age and argued that both of them – known from large numbers of isolated skull bones – were iguanids. Yet, *Anolbanolis* has never, to the best of my knowledge, been included in any phylogenetic analysis; and Conrad (2015), not mentioning *Anolbanolis* and not cited by Scarpetta (2020), had found the phylogenetic position of *Suzanniwana* difficult to resolve in the analysis of a dataset that included a much larger sample of early pan-iguanians.

Smith (2009: 312–313), incidentally, did not advertise *Suzanniwana* and *Anolbanolis* as the oldest iguanids either, accepting instead at least some of the even older jaw fragments that had been described as iguanid as “surely iguanid”, explicitly so for the “highly streamworn” over-62-Ma-old *Swainiguanoides* which had been described as “the oldest North American iguanid” (Sullivan, 1982). All of that and more was considered too uncertain by DeMar et al. (2017: 4, file S1: 26–28), who pointed out not only how fragmentary that material was (and that some of the Cretaceous specimens more likely belong to certain other squamate clades), but also that the presence of exclusive synapomorphies with Iguanidae (if confirmed) does not mean the specimens are actually inside that crown clade – they could be on its stem. As the “oldest definitive” iguanids, DeMar et al. (2017: 4) accepted *Anolbanolis*, followed by the uncontroversial *Afairiguana* which is younger than *Kopidosaurus*; curiously, they did not mention *Suzanniwana* at all.

The conclusion that the status of *Suzanniwana* and *Anolbanolis* (let alone *Swainiguanoides* and the like) is too uncertain and that *Kopidosaurus*, nowhere advertised for that purpose, should be used to set the minimum age for node 132 was accessible to me as an outsider to the fossil record of iguanians (or indeed squamates in general), but it took me several days of searching and reading papers and their supplementary information.

It took me much less effort to find that, under some phylogenetic hypotheses, the oldest known tetrapod (Materials and methods: Calibrations: Node 105 – Tetrapoda) is *Casineria*, a specimen I have studied in person and published on (Marjanović and Laurin, 2019); yet, the idea had never occurred to me or apparently anyone else in the field, even though its possibility should have been evident since 2017 and even though the phylogenetic hypotheses in question are by no means outlandish – one of them is even majoritarian.

In short, the paleontological literature is not optimized for divergence dating; the questions of which is the oldest known member of a group or when exactly that group evolved often take a back seat to understanding the anatomy, biomechanics, ecology, extinction, phylogeny or generally speaking evolution of that group in the minds of paleontologists – paleobiologists –, and this is reflected in the literature. Mining it for bounds on divergence dates is still possible, as I hope to have shown, but also rather exhausting.

## 3 Summary and conclusions

Irisarri et al. (2017) published the largest vertebrate timetree to date, calibrated with 30 minimum and 19 maximum ages for selected nodes (although one of each was not operational because the calibrations of other nodes set tighter constraints). With just three years of hindsight, only one of these dates stands up to scrutiny. Of the remaining 29 minimum ages, two had to be removed altogether, two had to be moved to previously uncalibrated nodes (with modifications to their numeric values), 15 were 4 Ma to 100 Ma too young, and 13 were 1 Ma to 96 Ma too old. Of the 19 maximum ages, seven had to be canceled altogether, while six were too young by 13 to 125 Ma and five too old by 3 to 93 Ma.

One of the minimum ages was taken from the wrong node in the cited secondary source, an earlier divergence-date analysis of molecular data (Noonan and Chippindale, 2006); another from the same source had a hundred million years added without explanation, most likely by typographic error. Only six of the 30 calibrated nodes were calibrated from primary literature. The calibration dates for seven nodes were taken from the compilation by Benton and Donoghue (2007), several from other compendia, four from Noonan and Chippindale (2006) who did not succeed in presenting the contemporary state of knowledge either.

Using software that was only able to treat all bounds as hard or all as soft (meaning that 2.5% or 5% of the credibility interval of each inferred node age must extend beyond the bound – younger than the minimum and older than the maximum age, where present), Irisarri et al. (2017) opted to treat all bounds as soft. For all minimum ages except one, this decision is not reproducible; it is even arguable for some of the maxima. This is not a purely theoretical problem; even the inferred mean ages of some calibrated nodes were younger than their minima in Marjanović (2019).

Redating of the tree of Irisarri et al. (2017) with the presumably improved calibrations results in many changes to the mean ages of nodes and to the sizes of their credibility intervals; not all of these changes are easily predictable.

Of the 276 references I have used to improve the calibrations, 50 were published in 2019, half of the total were published after mid-2016 (when Irisarri et al. seem to have completed the work on their manuscript), and 90% were published after mid-2005. Paleontology is a fast-moving field; secondary sources cannot keep up with the half-life of knowledge. A continually updated online compendium of calibration dates would be very useful, but the only attempt to create one (Ksepka et al., 2015) is no longer funded, has not been updated since early 2018, and had limited coverage. For the time being, each new attempt to calibrate node or tip ages will have to involve finding and studying the recent paleontological and chronostratigraphic literature on the taxa, strata and sites in question; although the Internet has made this orders of magnitude easier, it remains labor-intensive, in part because the the oldest record of a clade is often not published as such, but has to be inferred from comparing several sources on phylogeny, chronostratigraphy and sometimes taphonomy or even phylogenetics, as I illustrate here.

I urge that such work be undertaken and sufficiently funded. Accurate and precise timetrees remain an essential component of our understanding of, for example, the model organisms that are used in biomedical research: how much they can tell us about ourselves depends on how much evolution has happened along both branches since our last common ancestor, and that is in part a function of time.

## 5 Author Contributions

D. M. designed the experiments, gathered the data, interpreted the results, prepared the figure and the tables and wrote the paper.

## 8 Acknowledgments

Glory to our pirate queen, without whose work this paper would at best have taken a lot longer to write and at worst would have been severely outdated before submission.

Thanks to Albert Chen, Matteo Belvedere and Jason Pardo for an electronic reprint that would likely have been impossible to acquire in a timely manner otherwise; to Johan Renaudie for making me aware of another; to Olga Karicheva and the editorial office for several deadline extensions; to Jason Silviria and Paige dePolo for discussion of early eutherians; to the editor, Denis Baurain, for finding five reviewers; to all five reviewers and the editor for helpful comments; and to the editor and the editorial office for several more deadline extensions.

PhyloBayes only runs on Unix systems; Johan Renaudie (Museum für Naturkunde, Berlin) has access to such and kindly performed the analyses after expertly overcoming the gaps in the documentation of PhyloBayes.

